# A deletion in the *STA1* promoter determines maltotriose and starch utilization in *STA1*+ *Saccharomyces cerevisiae* strains

**DOI:** 10.1101/654681

**Authors:** Kristoffer Krogerus, Frederico Magalhães, Joosu Kuivanen, Brian Gibson

## Abstract

Diastatic strains of *Saccharomyces cerevisiae* are common contaminants in beer fermentations and are capable of producing an extracellular *STA1-*encoded glucoamylase. Recent studies have revealed variable diastatic ability in strains tested positive for *STA1,* and here we elucidate genetic determinants behind this variation. We show that poorly diastatic strains have a 1162 bp deletion in the promoter of *STA1*. With CRISPR/Cas9-aided reverse engineering, we show that this deletion greatly decreases the ability to grow in beer and consume dextrin, and the expression of *STA1*. New PCR primers were designed for differentiation of highly and poorly diastatic strains based on the presence of the deletion in the *STA1* promoter. In addition, using publically available whole genome sequence data, we show that the *STA1* gene is prevalent in among the ‘Beer 2’/’Mosaic Beer’ brewing strains. These strains utilize maltotriose efficiently, but the mechanisms for this have been unknown. By deleting *STA1* from a number of highly diastatic strains, we show here that extracellular hydrolysis of maltotriose through *STA1* appears to be the dominant mechanism enabling maltotriose use during wort fermentation in *STA1+* strains. The formation and retention of *STA1* seems to be an alternative evolutionary strategy for efficient utilization of sugars present in brewer’s wort. The results of this study allow for the improved reliability of molecular detection methods for diastatic contaminants in beer, and can be exploited for strain development where maltotriose use is desired.

## Introduction

Diastatic strains of *Saccharomyces cerevisiae* are considered spoilage microorganisms in industrial beer fermentations (Andrews and Gilliland, 1952; Meier-Dörnberg et al., 2018). These strains, formerly known as *Saccharomyces diastaticus*, are capable of producing an extracellular glucoamylase that enables fermentation of starch and oligosaccharides (Andrews and Gilliland, 1952; Erratt and Stewart, 1981; Kleinman et al., 1988; Yamashita et al., 1984). This in turn, causes super-attenuation in the beer, which leads to increased carbon dioxide and ethanol levels, as well as the possibility of off-flavours and a drier mouthfeel (Meier-Dörnberg et al., 2018). In the case of contamination in packaged beer, diastatic *S. cerevisiae* may even cause exploding bottles and cans, endangering the consumer.

The extracellular glucoamylase in diastatic *S. cerevisiae* is coded for by the *STA* genes (Tamaki, 1978; Yamashita et al., 1985b, 1985a). Three highly homologous *STA* genes (*STA1-3*) have been described (Lambrechts et al., 1991; Tamaki, 1978). In addition, *DEX1*-*2* has been used to describe genes encoding extracellular glucoamylases (Erratt and Stewart, 1981), however, these were later shown to be allelic to the *STA* genes (Erratt and Nasim, 1986). The *STA1* gene appears to be chimeric, consisting of rearranged gene fragments from both *FLO11* and *SGA1* (Lo and Dranginis, 1996; Yamashita et al., 1987). The 5’ end of *STA1* is homologous to *FLO11,* a gene encoding a membrane-bound flocculin promoting flocculation, while the 3’ end is homologous to *SGA1*, a gene encoding an intracellular glucoamylase used during sporulation. The catalytic domain is located in the *SGA1-*derived peptide, while the *FLO11-*derived peptide allows for extracellular secretion of the protein (Adam et al., 2004). The upstream sequences of *STA1* and *FLO11* are also nearly identical, meaning that these genes are largely co-regulated (Gagiano et al., 1999).

Current detection methods for diastatic *S. cerevisiae* rely mainly on either the microbiological detection through culturing on specialized selective growth media, or the molecular detection of the *STA1* gene through conventional or quantitative PCR (Brandl, 2006; Meier-Dörnberg et al., 2018; van der Aa Kühle, 1998; Yamauchi et al., 1998). The main weakness of the microbiological methods is that they are time-consuming. While the molecular methods are rapid, a recent study (Meier-Dörnberg et al., 2018) has revealed that there is considerable variability in diastatic ability and beer-spoilage potential in strains carrying the *STA1* gene. In fact, some strains that carry the *STA1* gene do not show spoilage potential, and these benign strains would be erroneously flagged as diastatic with the current molecular methods.

In this study, we examined the diastatic ability of a range of *STA1*+ *S. cerevisiae* strains. We also sequenced the open reading frame and upstream sequence of *STA1* in these same strains to search for polymorphisms that might explain the variable diastatic ability. Sequencing revealed that the poorly diastatic strains have a common 1162 bp deletion in the promoter of *STA1.* Through CRISPR/Cas9-aided reverse engineering, we show that this deletion greatly decreases diastatic ability and the expression of *STA1.* We designed new PCR primers targeting this deleted region, and these can be used to differentiate highly and poorly diastatic strains.

In addition, using publically available whole genome sequence data, we show that the *STA1* gene is prevalent in both the ‘Beer 2’ (’Mosaic Beer’) population, and surprisingly, the ‘French Guaina, human isolate’ population (Gallone et al., 2016; Peter et al., 2018). Strains in the ‘Beer 2’ population have been shown to utilize maltotriose efficiently, despite carrying frameshift mutations in *AGT1/MAL11* (Gallone et al., 2016). These strains are therefore likely to utilize alternative mechanisms for maltotriose use. By deleting *STA1* from a number of highly diastatic strains, we show here that *STA1* appears to have a central role in enabling maltotriose use in these strains during wort fermentation.

## Materials & Methods

### Yeast strains

A list of strains used in this study can be found in Table 1.

**Table 1.** Yeast strains used in the study.

| Code | Alternative name | Species | *STA1* | 1162 bp deletion in *STA1* promoter | Source |
| --- | --- | --- | --- | --- | --- |
| A15 | VTT-A83015 | *S. pastorianus* | - | NA | VTT Culture Collection |
| A60 | VTT-A75060 | *S. cerevisiae* | - | NA | VTT Culture Collection |
| WLP007 |  | *S. cerevisiae* | - | NA | White Labs Inc. |
| WLP023 |  | *S. cerevisiae* | - | NA | White Labs Inc. |
| TUM PI BA 109 |  | *S. cerevisiae* | + | - | BLQ Weihenstephan |
| TUM 3-D-2 |  | *S. cerevisiae* | + | - | BLQ Weihenstephan |
| TUM PI BA 45 |  | *S. cerevisiae* | + | + | BLQ Weihenstephan |
| TUM PI BB 105 |  | *S. cerevisiae* | + | + | BLQ Weihenstephan |
| TUM PI BA 31 |  | *S. cerevisiae* | + | + | BLQ Weihenstephan |
| TUM 71 |  | *S. cerevisiae* | + | - | BLQ Weihenstephan |
| TUM 3-H-2 |  | *S. cerevisiae* | + | - | BLQ Weihenstephan |
| WY3711 |  | *S. cerevisiae* | + | - | Wyeast Laboratories |
| WLP565 |  | *S. cerevisiae* | + | + | White Labs Inc. |
| WLP566 |  | *S. cerevisiae* | + | + | White Labs Inc. |
| WLP570 |  | *S. cerevisiae* | + | + | White Labs Inc. |
| WLP644 |  | *S. cerevisiae* | + | + | White Labs Inc. |
| Ontario Farmhouse |  | *S. cerevisiae* | + | + | Escarpment Laboratories |
| Wild Thing |  | *S. cerevisiae* | + | + | Escarpment Laboratories |
| A62 | VTT-A81062 | *S. cerevisiae* | + | + | VTT Culture Collection |
| **CRISPR/Cas9-modified strains** |  |  |  |  |  |
| WY3711_D1 | P<sub>sta1</sub>-Δ1 (*STA1*:c.-1370_-290del) | *S. cerevisiae* | + | + | This study |
| WY3711_S1 | *sta1*-Δ0 (*STA1*:c.-1370_+2421del) | *S. cerevisiae* | - | NA | This study |
| TUM PI BA 109_S1 | *sta1*-Δ0 (as above) | *S. cerevisiae* | - | NA | This study |
| TUM 71_S1 | *sta1*-Δ0 (as above) | *S. cerevisiae* | - | NA | This study |

### Dextrin fermentation

The ability to ferment dextrin was tested in minimal growth media with dextrin as the sole carbon source. Strains were grown overnight in YP-Glucose (2%), after which 20 µL of overnight culture was used to inoculate 2 mL microcentrifuge tubes containing 1 mL of dextrin media (0.67% YNB without amino acids, 1% dextrin from potato starch). The tubes were incubated at room temperature for 3 weeks. Samples were drawn each week, and analysed with HPLC to determine the amount of dextrin consumed.

### Growth on starch agar

The ability to grow on solid media containing starch as the sole carbon source was tested on agar plates (Meier-Dörnberg et al., 2018). Agar plates were prepared containing 0.67% YNB /wo amino acids, 1.5% soluble starch (Merck, Germany) and 40 mg/L bromophenol blue. pH was adjusted to 5.2 with 0.1M HCl. Yeast strains were grown overnight in YP-Glucose, washed twice with sterile MQ-H_2_O, and then resuspended to an OD600 of 1. Aliquots of yeast suspension (100 µL) were spread out on agar plates. The plates were incubated in an anaerobic jar (Anoxomat AN2CTS, Mart Microbiology) at 25 °C for four weeks.

### Growth in beer

The spoilage potential of *STA1*+ strains was assessed by growing them in beer. Beer was produced by inoculating one litre of 10 °Plato all-malt wort with *S. pastorianus* A15 and fermenting at 25 °C for one week. The beer was centrifuged and sterile-filtered (0.45 µm, followed by 0.22 µm). The beer was analysed with HPLC and determined to contain less than 1 g/L each of maltose and maltotriose. Yeast strains were grown overnight in YP-Glucose, washed twice with sterile MQ-H_2_O, and then resuspended to an OD600 of 1. Sterile-filtered beer (4750 µL) was inoculated with 250 µL of pre-culture for a starting OD600 of 0.05. The beer cultivations were incubated statically in an anaerobic jar (Anoxomat AN2CTS, Mart Microbiology) at 25 °C for three weeks. The optical density was monitored once a week, and the final amount of dextrin consumed was analysed with HPLC after three weeks.

### Wort fermentation and analysis

80mL-scale fermentations were carried out in 100mL Schott bottles capped with glycerol-filled airlocks. Yeast strains were grown overnight in 25 mL YP-Maltose (4%) at 25 °C. The pre-cultured yeast was then inoculated into 80 mL of 15 °P all-malt wort at a rate of 1 g fresh yeast L*^-^*^1^. Fermentations were carried out in triplicate at 25 °C for 15 days. Fermentations were monitored by mass lost as CO_2_. Samples were also drawn throughout fermentation and analysed with HPLC to monitor sugar consumption.

Concentrations of fermentable sugars and dextrin were measured by HPLC using a Waters 2695 Separation Module and Waters System Interphase Module liquid chromatograph coupled with a Waters 2414 differential refractometer (Waters Co., Milford, MA, USA). An Aminex HPX-87H organic acid analysis column (300 × 7.8 mm, Bio-Rad) was equilibrated with 5 mM H_2_SO_4_ (Titrisol, Merck, Germany) in water at 55 °C and samples were eluted with 5 mM H_2_SO_4_ in water at a 0.3 ml/min flow rate.

### Maltotriose uptake assay

The zero-trans maltotriose uptake rate was assayed using [U-^14^C]-maltotriose as described by Lucero et al. (1997). Yeast strains were grown in YP-Maltose at 20 °C to an OD600 between 4 and 8 prior to uptake measurement. The uptake rate was determined at 20 °C using 5 mM of [U-^14^C]-maltotriose with 1 min incubation time. [U-^14^C]-maltotriose was repurified before use as described by Dietvorst et al. (2005).

### Sequencing of STA1

The *STA1* open reading frame and promoter (2.5 kb upstream) were amplified by PCR and then sequenced using Sanger sequencing. A nested PCR approach was used to prevent PCR primers from amplifying fragments from *FLO11* and *SGA1*. First, a 5.2 kb amplicon was produced using primers STA1_Full_Fw and STA1_Full_Rv (Table 2). The amplicon was diluted 1:500 in MQ-H_2_O and used as template DNA for the ten PCR reactions described in Table 2. All PCR reactions were carried out with Phusion High-Fidelity PCR Master Mix with HF Buffer (Thermo Scientific) and primer concentrations of 0.5 µM. Amplicons were cleaned using the QIAquick PCR purification kit of Qiagen (Hilden, Germany), and sequenced at Seqlab-Microsynth (Goettingen, Germany). Sequences were aligned in Geneious 10.0.9 (Biomatters). The sequences are available in the Supplementary material.

**Table 2.**
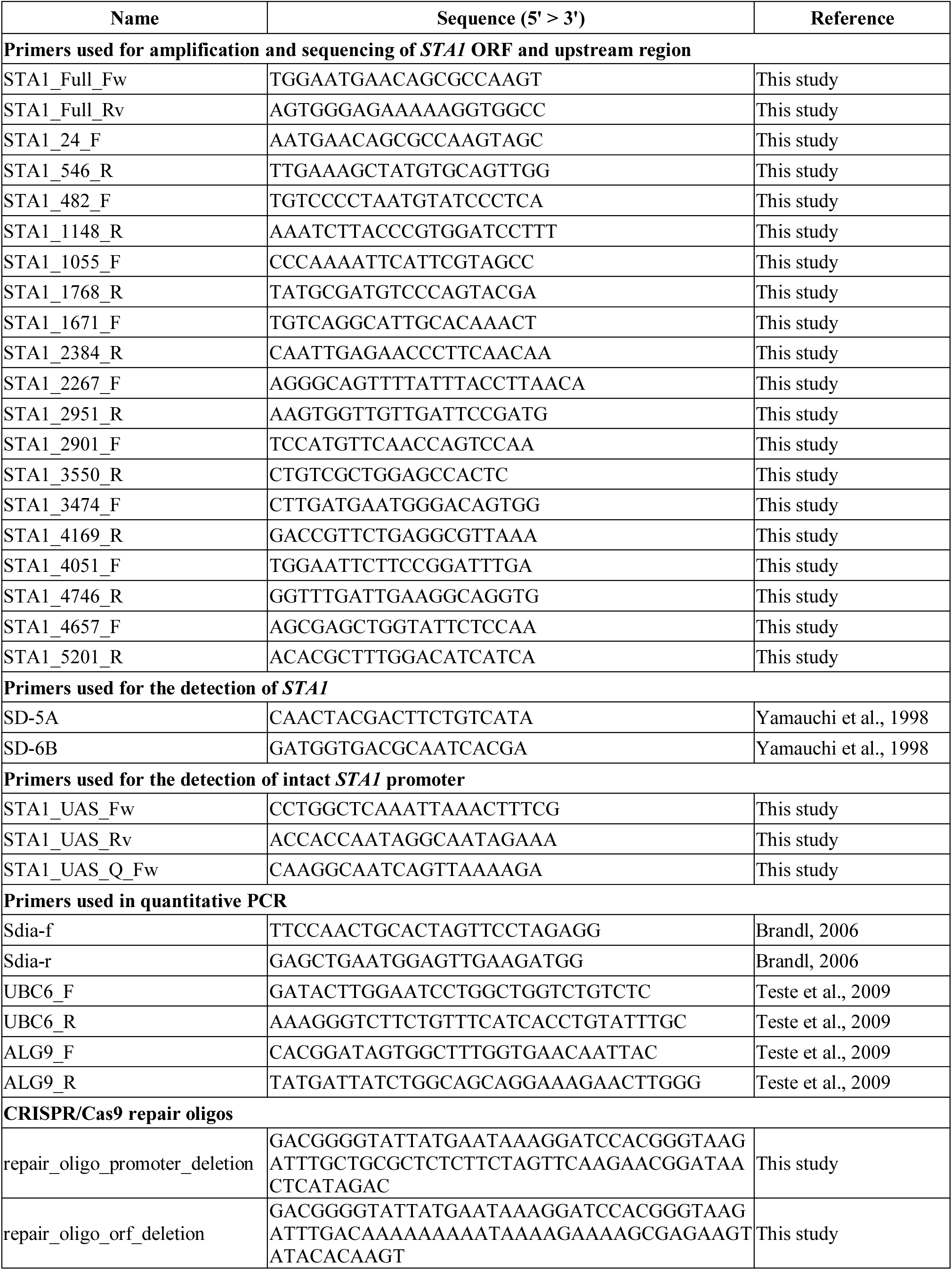
Oligonucleotides used in the study.

### Multiplex PCR to detect presence of deletion in STA1 promoter

The presence of the *STA1* gene was tested with PCR using the previously published primers SD-5A and SD-6B (Yamauchi et al., 1998). These primers amplify an 868 bp fragment from the *STA1* gene. In addition, a new primer pair (STA1_UAS_Fw and STA1_UAS_Rv; Table 2) was designed to detect whether the *STA1+* yeast strains had the full promoter sequence. Both primers bind within the 1162 bp region that was found to be deleted in the poorly diastatic strains, and they amplify a 599 bp fragment. Primers were tested in separate and multiplex PCR reactions. PCR reactions were carried out with Phusion High-Fidelity PCR Master Mix with HF Buffer (Thermo Scientific) and primer concentrations of 0.5 µM. The following PCR program was used: 98 °C 30 sec, (98 °C 10 sec, 63 °C 15 sec, 72 °C 30 sec) × 30 cycles, 72 °C 10 min. PCR products were separated and visualized on 1.0% agarose gels.

To test the primers in a simulated real-world scenario, we performed PCR on DNA extracted from cultures of lager yeast contaminated with various ratios of diastatic *S. cerevisiae*. Cultures of *S. cerevisiae* WY3711 and *S. pastorianus* A15 were grown overnight in YP-Glucose, and duplicate aliquots of 3 · 10^7^ cells containing 10^-1^ to 10^-6^ fractions of *S. cerevisiae* WY3711 were prepared in ten-fold dilutions with *S. pastorianus* A15. Aliquots of *S. cerevisiae* WY3711 and *S. pastorianus* A15 were prepared as positive and negative controls, respectively. DNA was extracted from these aliquots using a YeaStar Genomic DNA kit (Zymo Research, USA), and PCR was performed on 50 ng template DNA as described above.

### Quantitative PCR to detect presence of deletion in STA1 promoter

The presence of the *STA1* gene was tested with quantitative PCR using the previously published primers Sdia-f and Sdia-r (Brandl, 2006). These primers amplify a 79 bp fragment from the *STA1* open reading frame. In addition, a new primer pair (STA1_UAS_Q_Fw and STA1_UAS_Rv; Table 2) was designed to detect whether the *STA1+* yeast strains had the full promoter sequence. Both primers bind within the 1162 bp region that was found to be deleted in the poorly diastatic strains, and amplify a 223 bp fragment. The qPCR reactions were prepared with PerfeCTa SYBR® Green SuperMix (QuantaBio, USA) and 0.3 µM of the primers (Table 2). The qPCR reactions were performed on a LightCycler® 480 II instrument (Roche Diagnostics, Switzerland) in two technical replicates on 50 ng template DNA extracted with a YeaStar Genomic DNA kit (Zymo Research, USA). The following program was used: pre-incubation (95 °C for 3 min), amplification cycle repeated 45 times (95 °C for 15 s, 60 °C for 30 s, 72 °C for 20 s with a single fluorescence measurement), melting curve program (65–97 °C with continuous fluorescence measurement), and finally a cooling step to 40 °C.

### CRISPR/Cas9 mediated deletions

In order to confirm the effect of the 1162 bp deletion that was observed in the *STA1* promoter of the poorly diastatic strains, reverse engineering in the highly diastatic *S. cerevisiae* WY3711 strain was performed using the CRISPR/Cas9 system. Plasmid construction was carried out using the plasmid pCC-036 as backbone (Rantasalo et al., 2018). pCC-036 contains yeast codon-optimized Cas9 expressed under *TDH3p*, guiding RNA (gRNA) expressed under *SNR52p*, and *hygR* for selection on hygromycin. The gRNA protospacer sequence, TGGCTCAAATTAAACTTTCG, was designed using CCTop (Stemmer et al., 2015). The protospacer sequence was designed to induce a double-stranded break within the region to be deleted and was chosen for minimal off-target activity. A synthetic DNA fragment with the gRNA sequence was ordered from Integrated DNA Technologies (Belgium) as a gBlock, and introduced into the plasmid with restriction enzyme-based techniques (Thermo Scientific, Finland). The ligated plasmid was transformed into *E. coli* TOP10 by electroporation, and plasmid correctness was confirmed by Sanger sequencing. A 80 bp repair oligo (repair_oligo_promoter_deletion), consisting of adjacent 40 bp sequences homologous to those up- and downstream of the deleted region (Table 2), was also ordered from Integrated DNA Technologies (Belgium).

Transformation of *S. cerevisiae* WY3711 was performed using a standard lithium acetate-based protocol with 40 minute incubation at 42 °C (Gietz and Woods, 2002). Cells were transformed together with 3.6 µg of purified plasmid and 2.5 nmol of repair oligo (double-stranded). The transformed cells were selected on plates containing 300 mg/L Hygromycin B (Sigma-Aldrich). Colony PCR using primer pairs 1055F/2951R and STA1_UAS_Fw/STA1_UAS_Rv, and Sanger sequencing, was used to confirm successful deletion in the transformed cells. Colonies from selection plates were replated three times onto YPD agar plates to encourage plasmid loss, after which they were stored at −80°C.

The same CRISPR/Cas9 plasmid that was used above to delete the 1162 bp region in the *STA1* promoter, was also used to delete the entire *STA1* open reading frame from three of the highly diastatic *S. cerevisiae* strains (TUM PI BA 109, TUM 71 and WY3711). A 80 bp repair oligo (repair_oligo_orf_deletion; Table 2), consisting of adjacent 40 bp sequences homologous to those up- and downstream of the region to be deleted (−1370 to +2421) was transformed together with the purified plasmid as described above. Colony PCR using primer pairs STA1_Full_Fw/STA1_Full_Rv, 1055F/5201R and SD-5A/SD-6B, and Sanger sequencing, was used to confirm successful deletion in the transformed cells (Supplementary Figure S4).

### STA1 transcript analysis by RT-qPCR

Strains were grown overnight in YP-Glucose, after which four replicate cultures per strain were started by inoculating 20 mL YPGE (1% yeast extract, 2% peptone, 3% glycerol and 2% ethanol) to a starting OD600 of 0.1. Cultures were overnight at 25 °C (OD600 varied between 2 and 4), after which RNA was isolated from pelleted yeast using hot formamide extraction (Shedlovskiy et al., 2017). RNA was precipitated by first diluting 50 µL RNA solution with 25 µL 3M sodium acetate and 200 µL nuclease-free water, after which 825 µL ice-cold ethanol was added and solutions were stored overnight at −20 °C. The solutions were centrifuged at 16000 × *g* for 30 minutes, after which the pellet was washed with ice-cold 75% ethanol and tubes were centrifuged again at 8000 × *g* for 5 minutes. The supernatant was removed and the pellet was allowed to air-dry for up to 30 minutes, after which the RNA pellet was dissolved in nuclease-free water. The RNA solution was treated with TURBO DNAse (DNA-Free kit, Invitrogen) and the DNAse was subsequently inactivated using the supplied inactivation reagent. RNA was quantified with a Qubit 2.0 fluorometer and its quality was assessed on 1.2% agarose gels (made in 1×TAE buffer).

250 ng of total RNA was reverse-transcribed using a qScript Flex cDNA kit (QuantaBio, USA), using a mixture of supplied oligo-dT and random primers according to kit instructions. The resulting cDNA was diluted 5-fold and 4 µL of diluted cDNA (corresponding to 10 ng of total RNA) was used as template in 20 µL qPCR reactions. The qPCR reactions were prepared with PerfeCTa SYBR® Green SuperMix (QuantaBio, USA) and 0.3 µM of gene-specific primers (Table 2). In addition to using primers specific to *STA1,* reactions with primers for two house-keeping genes (*ALG9* and *UBC6*) were also performed (Teste et al., 2009). The qPCR reactions were performed on a LightCycler® 480 II instrument (Roche Diagnostics, Switzerland) in two technical replicates on the reverse-transcribed RNA isolated from four biological replicates. The following program was used: pre-incubation (95 °C for 3 min), amplification cycle repeated 45 times (95 °C for 15 s, 60 °C for 30 s, 72 °C for 20 s with a single fluorescence measurement), melting curve program (65–97 °C with continuous fluorescence measurement), and finally a cooling step to 40 °C. The relative expression of *STA1* in the three examined yeast strains was calculated using the ‘delta-delta C_T_’-method by normalizing expression to that of the two house-keeping genes *ALG9* and *UBC6* (Pfaffl, 2001).

### Oxford Nanopore MinION whole genome sequencing

The genomes of *S. cerevisiae* WY3711 and A62 were sequenced using an Oxford Nanopore MinION. Genomic DNA was extracted with a YeaStar Genomic DNA kit (Zymo Research, USA) and then barcoded with the Native Barcoding kit (EXP-NBD104; Oxford Nanopore Technology, United Kingdom). A 1D sequencing library was then prepared from the barcoded DNA using the Ligation Sequencing Kit (SQK-LSK109; Oxford Nanopore Technology, United Kingdom). The library was sequenced on a FLO-MIN106D (R9.4.1) flow cell with a MinION device (Oxford Nanopore Technology, United Kingdom). The raw fast5 files were basecalled using the GPU-version of Guppy (version 2.3.7; using the supplied dna_r9.4.1_450bps_flipflop.cfg configuration). The basecalled reads were demultiplexed with qcat (version 1.0.1; available from https://github.com/nanoporetech/qcat), and filtered to a minimum length of 1,000 bp and average read quality score of 10 using NanoFilt (version 2.2.0; available from https://github.com/wdecoster/nanofilt). This resulted in a total of 1.3 Gbp of sequence for *S. cerevisiae* WY3711 (coverage of 108×) and 1.9 Gbp of sequence for *S. cerevisiae* A81062 (coverage of 155×).

### De novo assembly

The sequencing reads generated in this study (for *S. cerevisiae* WY3711 and *S. cerevisiae* A62) were *de novo* assembled using the LRSDAY (version 1.4) pipeline (Yue and Liti, 2018). The initial assemblies were produced with smartdenovo (available from https://github.com/ruanjue/smartdenovo) using default settings. The assemblies were then polished twice with Nanopolish (0.11.1; available from https://github.com/jts/nanopolish). Alignment of long reads was performed with minimap2 (version 2.16; Li, 2018). The contigs in the polished assemblies were then scaffolded with Ragout (version 2.0; Kolmogorov et al., 2014) to *S. cerevisiae* S288C (R64-2-1). Assembly statistics are available in Supplementary Table S5 and Supplementary Figure S5.

In addition, *de novo* assembly of long sequencing reads produced in other studies (accession numbers are available in Supplementary Table S1) was also carried out using the LRSDAY pipeline essentially as described above. The assemblies were polished twice with Pilon (version 1.22; Walker et al., 2014) using Illumina reads (accession numbers are available in Supplementary Table S1). Identity between assemblies was determined with ‘dnadiff’ from MUMmer (version 3.23; Kurtz et al., 2004).

### Prevalence of STA1 and phylogenetic analysis

In addition to the two genome assemblies produced from sequencing reads generated in this study, we retrieved publically available genome assemblies of *S. cerevisiae*. Genome assemblies of the 157 *S. cerevisiae* strains described in Gallone et al. (2016) were retrieved from NCBI (BioProject PRJNA323691). Genome assemblies of the 1011 *S. cerevisiae* strains described by Peter et al. (2018) were retrieved from https://www.yeastgenome.org/1011-yeast-genomes. The genome assembly of *S. cerevisiae* A81062 (Krogerus et al., 2016) was retrieved from NCBI (Assembly ASM193724v1). The prevalence of the *STA1* gene in natural *S. cerevisiae* isolates was investigated by performing a BLAST search of sequence ‘STA1_BLAST’ (Supplementary Table S2) in the collected genome assemblies using NCBI-BLAST (version 2.6.0).

Multiple sequence alignment of the *S. cerevisiae* assemblies was performed with the NASP pipeline (Roe et al., 2016) using *S. cerevisiae* S288C (R64-2-1) as the reference genome. A matrix of single nucleotide polymorphisms (SNP) in the 1171 strains was extracted from the aligned sequences. The SNPs were annotated with SnpEff (Cingolani et al., 2012) and filtered as follows: only sites that were in the coding sequence of genes, present in all 1171 strains and with a minor allele frequency greater than 0.25% were retained (in at least 3 strains). The filtered matrix contained 26 725 238 SNPs (462842 sites). A maximum likelihood phylogenetic tree was estimated using IQ-TREE (Nguyen et al., 2015). IQ-TREE was run using the ‘GTR+F+R4’ model and 1000 ultrafast bootstrap replicates (Minh et al., 2013). The resulting maximum likelihood tree was visualized in FigTree and rooted with the Taiwanese outgroup.

### Data visualization and analysis

Data and statistical analyses were performed with R (http://www.r-project.org/). Plots were produced in R and FigTree.

### Data availability

The Sanger sequences generated in this study are available in the Supplementary material, and the long sequencing reads have been submitted to NCBI-SRA under BioProject number PRJNA544899 in the NCBI BioProject database (https://www.ncbi.nlm.nih.gov/bioproject/).

## Results

### Screening of diastatic ability

The diastatic ability of eighteen *S. cerevisiae* and one *S. pastorianus* strain (Table 1) was tested using three different tests: growth in beer, growth on starch agar, and fermentation of dextrin as a sole carbon source. Of the eighteen *S. cerevisiae* strains, 15 tested positive for *STA1* using the SD-5A/SD-6B primer pair (Yamauchi et al., 1998). Despite carrying the *STA1* gene, only five out of 15 strains, were able to grow in beer and on starch agar (Figure 1A). These five strains also fermented dextrin efficiently. All four of the strains that tested negative for *STA1* were unable to grow in beer or on starch agar, and did not consume any dextrin.

**Figure 1.**
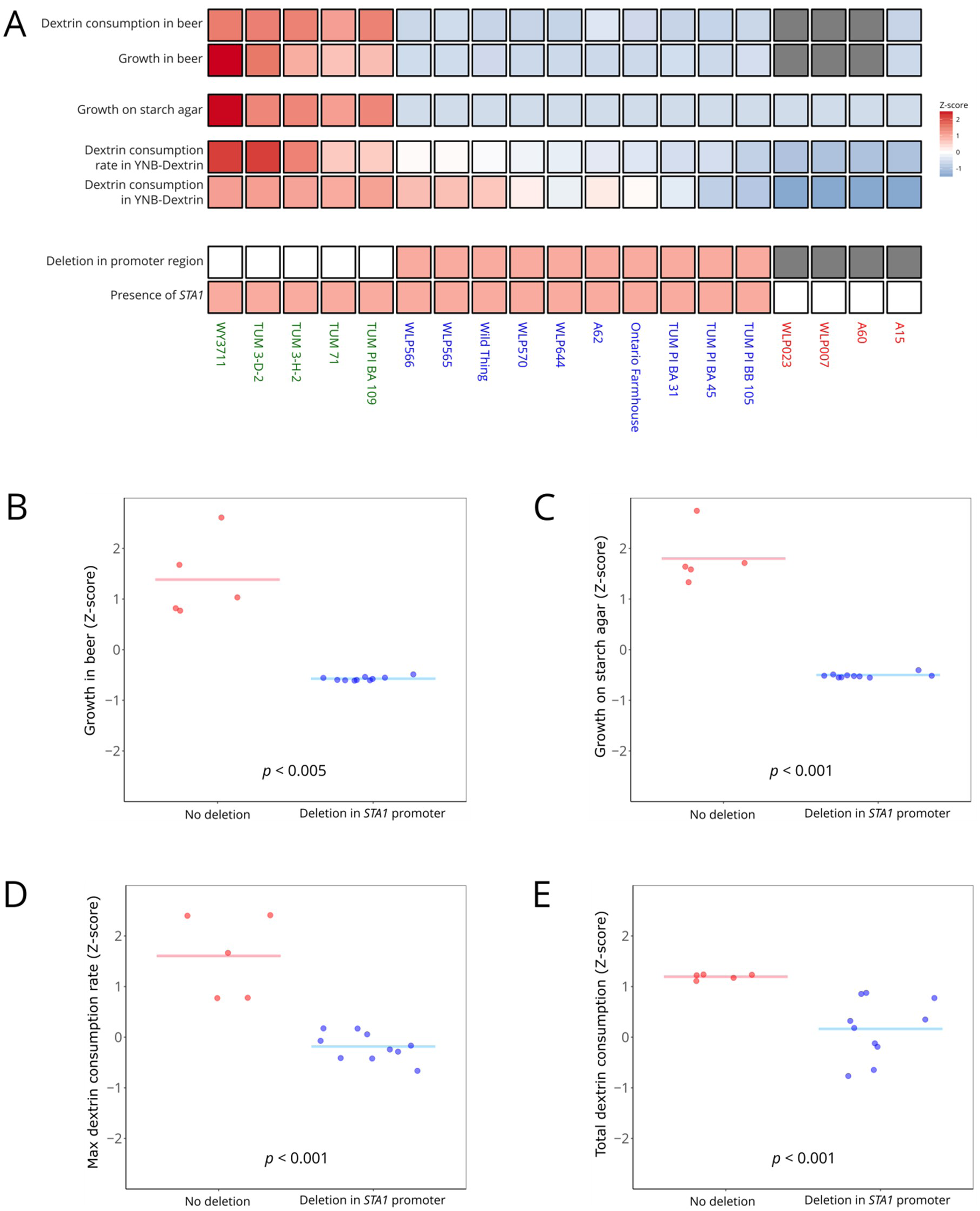
A comparison of the diastatic ability of 15 *STA1+* and 4 *STA1-* yeast strains. **(A)** A heatmap of the diastatic ability based on three different tests in the 19 *Saccharomyces* strains. The heatmap is coloured based on Z-scores (red and blue indicate values higher and lower than average, respectively). The strains are coloured as follows; green: *STA1*+, no deletion in *STA1* promoter; blue: *STA1+,* deletion in *STA1* promoter; red: *STA1-*. (**B-E**) Pairwise comparison of the results from the individual diastatic tests (**B:** growth in beer; **C:** growth on starch agar; **D-E:** fermentation of dextrin) between *STA1+* strains with no deletion in the *STA1* promoter compared to strains with a deletion in the *STA1* promoter. The statistical significance between the two groups was tested with the Mann-Whitney U test.

We then amplified and sequenced the *STA1* open reading frame and a 2.5kb upstream region, to search for polymorphisms that might explain the variation in diastatic ability in the 15 strains that tested positive for *STA1.* No nonsense mutations in the open reading frame of *STA1* were observed in any of the strains. However, we did observe a 1162 bp deletion upstream of *STA1* (−1370 to −209 from the start codon) in 10 out of the 15 strains (Figure 2 and Figure 1A). This 1162 bp region contains an upstream activation sequence and transcription factor (Ste12 and Tec1) binding site (Kim et al., 2004a, 2004b). Interestingly, the presence of this deletion appeared to coincide with decreased diastatic ability (Figure 1). The strains with the deletion in the promoter exhibited significantly less diastatic ability in all tests (Figure 1B-E). We hypothesised that this deletion impedes transcription of *STA1*, which in turn decreases the amount of *STA1-*derived glucoamylase produced by the strains.

**Figure 2.**
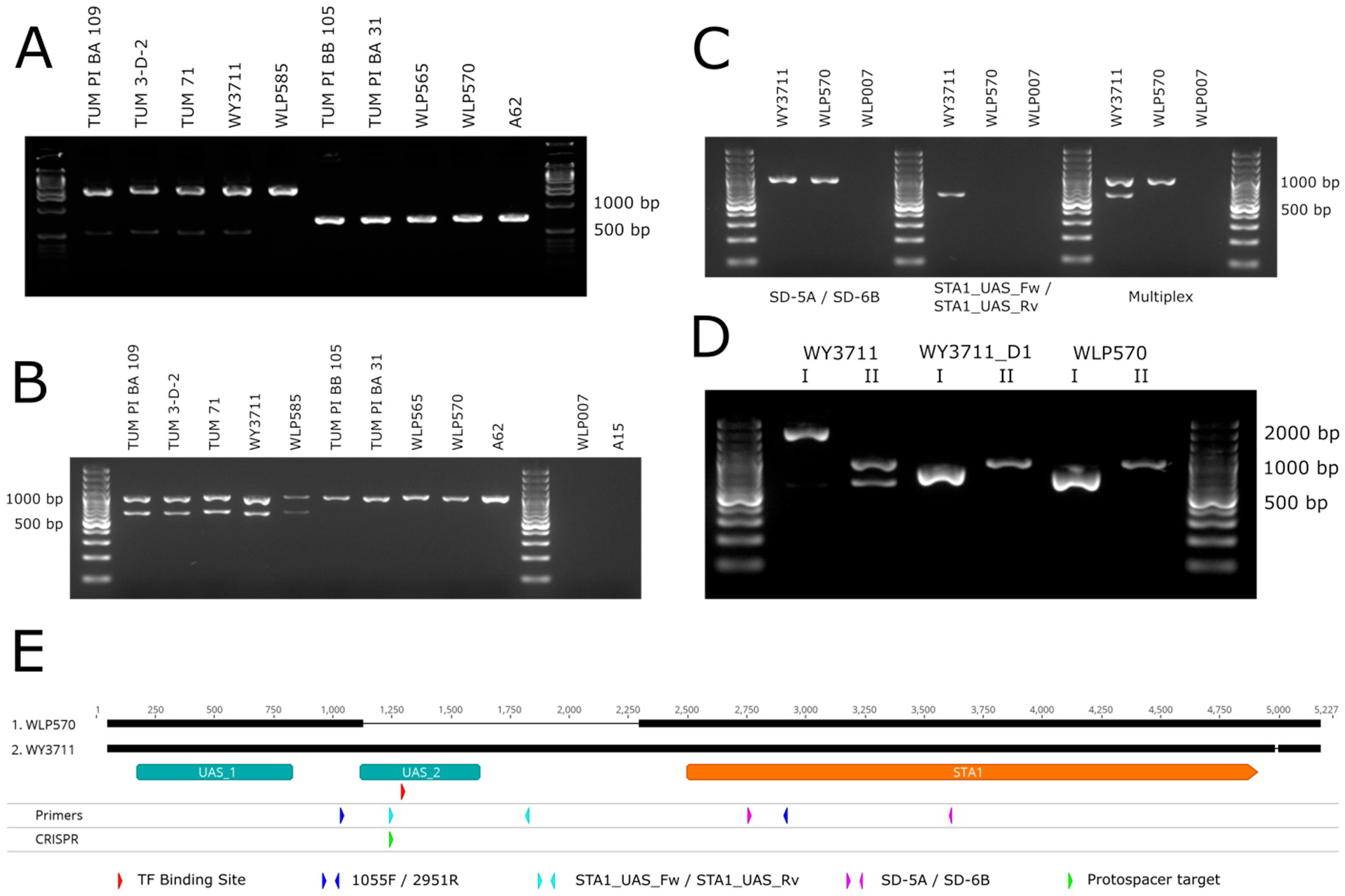
The 1162 bp deletion in the *STA1* promoter. **(A)** PCR products from 10 *STA1+* strains using primers STA1_1055_F/ STA1_2951_R. **(B)** Multiplex PCR products from the same 10 *STA1+* strains as **(A)** and two *STA1-* controls using primers SD-5A/SD-6B and STA1_UAS_Fw/STA1_UAS_Rv. **(C)** Individual and multiplex PCR reactions with WY3711 (*STA1+,* no deletion in *STA1* promoter), WLP570 (*STA1+,* deletion in *STA1* promoter), and WLP007 (*STA1-*) using primers SD-5A/SD-6B and STA1_UAS_Fw/STA1_UAS_Rv. (**D**) Confirmation of successful deletion of the 1162 bp region in the *STA1* promoter in strain WY3711_D1 using primers STA1_1055_F/STA1_2951_R (represented by I) and the multiplex primers SD-5A/SD-6B and STA1_UAS_Fw/STA1_UAS_Rv (represented by II). (**E**) Sequence alignment of *STA1* open reading frame and upstream region from WY3711 and WLP570, and locations of PCR primers, transcription factor binding site (TF) and CRISPR protospacer target. UAS_1 and UAS_2 are locations of upstream activation sequences described in Kim et al. (2004a, 2004b).

### Confirmation by reverse engineering

To confirm that the 1162 bp deletion that was observed in the *STA1* promoter had a negative effect on diastatic ability and *STA1* expression, we performed reverse engineering aided by the CRISPR/Cas9 system. The gRNA protospacer sequence was designed to cause a double-stranded break within the region to be deleted (Figure 2E). The 1162 bp region (−1370 to −209 upstream of the *STA1* open reading frame) was deleted from the highly diastatic WY3711 strain by transformation with a Cas9- and gRNA-expressing plasmid and an 80 bp double-stranded repair oligo. PCR using primer pairs binding within (STA1_UAS_Fw / STA1_UAS_Rv) and outside the deleted region (1055F / 2951R), together with Sanger sequencing, was used to confirm successful deletion (Figure 2D and Supplementary Figure S4).

The deletion strain, WY3711_D1, was then tested for diastatic ability. WY3711, the wild-type strain from which WY3711_D1 was derived, and WLP570, a strain that naturally contains the 1162 bp deletion in the promoter, were included as controls. In all tests, the diastatic ability of WY3711_D1 was less than WY3711. When inoculated into beer, WY3711_D1 and WLP570 grew significantly less than WY3711 (Figure 3A). In contrast to WY3711, WY3711_D1 and WLP570 had also not consumed any dextrin from the beer after 3 weeks (Figure 3D). When the strains were grown in minimal media with dextrin as the sole carbon source, we saw negligible dextrin consumption after 3 weeks with WY3711_D1 and WLP570 when cultures were incubated anaerobically (Figure 3B). Interestingly, we observed delayed activity in these strains when cultures were incubated aerobically (Figure 3C). WY3711 performed similarly in both anaerobic and aerobic conditions, and in both cases consumed significantly more dextrin than the strains with the deletion in the promoter.

**Figure 3.**
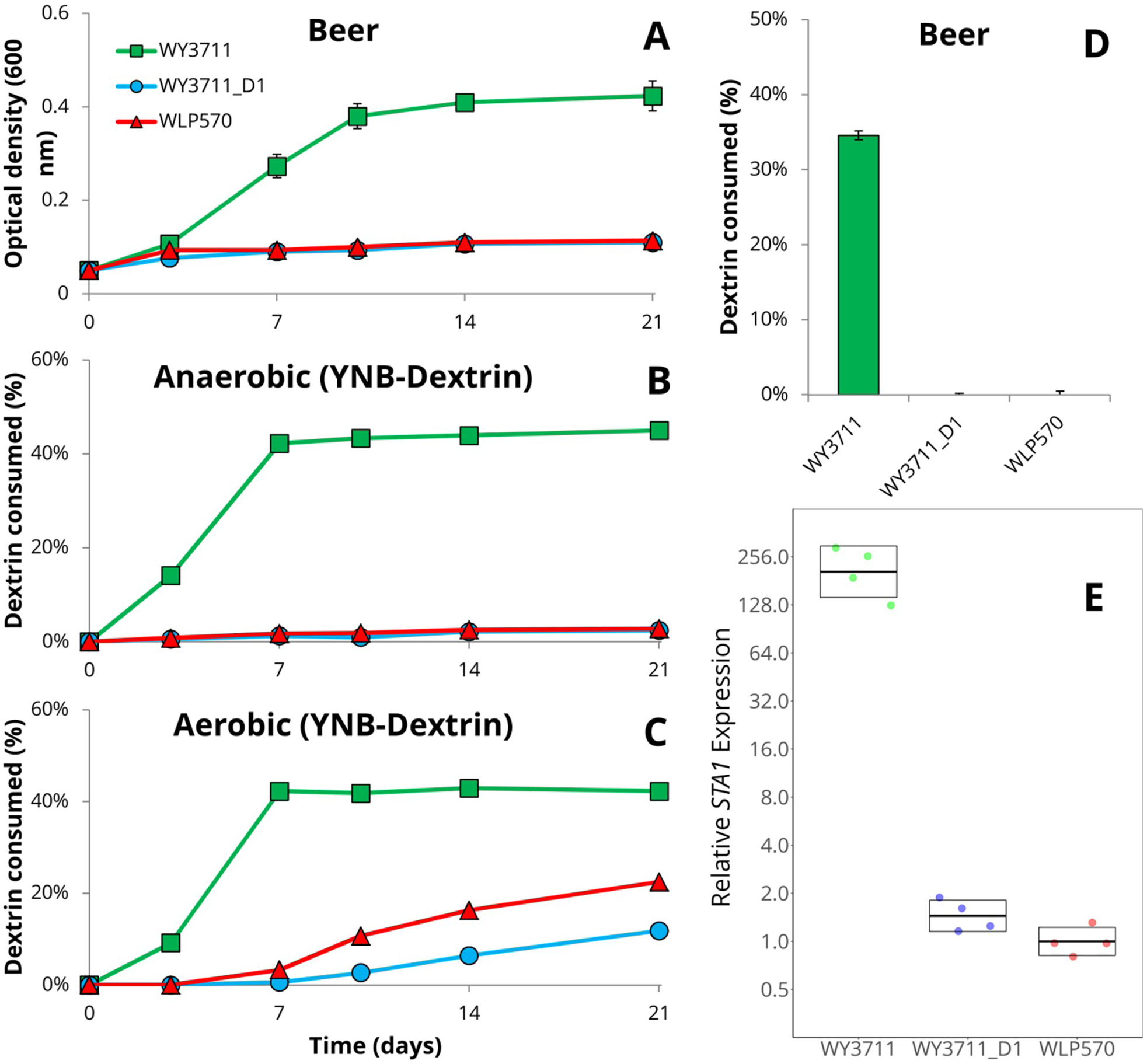
Confirmation of the role of the 1162 bp deletion in the *STA1* promoter by reverse engineering. Three *STA1+* strains were compared: WY3711 (no deletion in *STA1* promoter), WY3711_D1 (CRISPR-mediated deletion in *STA1* promoter), and WLP570 (natural deletion in *STA1* promoter). **(A)** The optical density (600 nm) when strains were inoculated into beer at a starting value of 0.1. Values for WY3711 were different from the two other strains starting from day 7 (*p* < 0.01 by two-tailed Student’s t-test). (**B**-**C**) The amount of dextrin consumed from YNB-Dextrin media in anaerobic (**B**) and aerobic (**C**) conditions. Values for WY3711 were different from the two other strains starting from day 4 (*p* < 0.01 by two-tailed Student’s t-test). (**D)** The amount of dextrin consumed from the beer (**A**) after 3 weeks of incubation. (**E**) The relative expression of *STA1* (normalized to *ALG9* and *UBC6*) determined by RT-qPCR in derepressed conditions. Points indicate values from four biological replicates and boxes indicate the mean and standard deviation. Values for WY3711 were significantly higher than those of the two other strains (*p* < 0.01 by two-tailed Student’s t-test).

To test the effect that the 1162 bp deletion in the *STA1* promoter has on the expression of *STA1,* we performed RT-qPCR analysis on RNA extracted from exponential phase cultures in YPGE. This growth medium was chosen as glucose represses *STA1* expression (Kim et al., 2004b; Pretorius et al., 1986). We observed a 150-200 fold higher abundance of *STA1* mRNA in the samples from WY3711 compared to WY3711_D1 and WLP570, respectively (Figure 3E). These results suggest that the decreased diastatic ability observed in the *STA1*+ strains with the 1162 bp deletion is a result of decreased expression of *STA1*.

### Designing primers for the detection of the full STA1 promoter

To improve the reliability of the molecular detection methods for diastatic *S. cerevisiae*, we designed two new primer pairs that bind within the 1162 bp region that is absent in the poorly diastatic *STA1+* strains. These new primers can therefore be used to differentiate whether a strain that has tested positive for *STA1* with the SD-5A / SD-6B primers (Yamauchi et al., 1998) is likely to be highly diastatic or not. The first primer pair (STA1_UAS_Fw / STA1_UAS_Rv) was designed to produce a 599 bp amplicon, and can be used together with the SD-5A / SD-6B primers in a single multiplex PCR reaction (Figure 2B-C). Here, strains with the full *STA1* promoter produce two amplicons (599 and 868 bp), while poorly diastatic strains only produce a single amplicon (868 bp). To test how this multiplex PCR reaction would perform in a simulated brewery scenario, we contaminated a lager yeast culture with varying ratios of diastatic *S. cerevisiae* WY3711 (10^-6^ – 10^-1^). We were able to detect the presence of diastatic *S. cerevisiae* at a concentration of 10^-4^ (Figure 4A).

**Figure 4.**
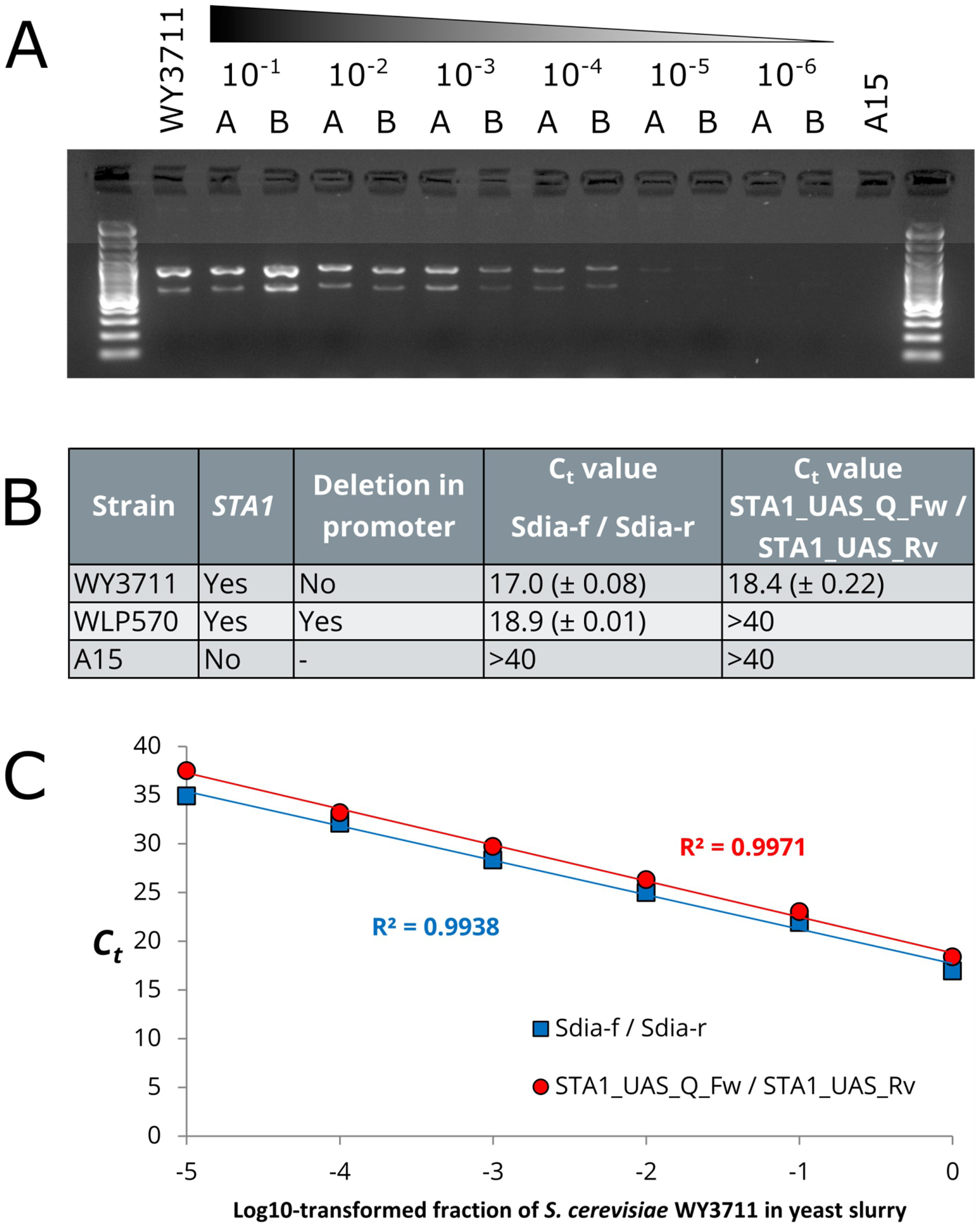
The sensitivity of the newly designed PCR primers for differentiation of *STA1+* strains with and without the 1162 bp deletion in the *STA1* promoter. **(A)** Multiplex PCR products using primers SD-5A/SD-6B and STA1_UAS_Fw/STA1_UAS_Rv from DNA extracted from duplicate *S. pastorianus* A15 cultures contaminated with increasing ten-fold ratios of contamination by *S. cerevisiae* WY3711 (values 10^-1^-10^-6^ indicate fraction of WY3711 in the culture). (**B-C**) The cycle threshold (C_t_) values of quantitative PCR reactions using DNA extracted as in (**A**) using primers STA1_UAS_Q_Fw/STA1_UAS_Rv and Sdia-f/Sdia-r.

In addition, a second primer pair (STA1_UAS_Q_Fw / STA1_UAS_Rv) was also designed to be useable in quantitative PCR reactions where a shorter amplicon is desired. This primer pair produces a 223 bp amplicon. We performed quantitative PCR reactions on DNA extracted from *S. cerevisiae* WY3711, *S. cerevisiae* WLP570 and *S. pastorianus* A15 using both the previously published Sdia-f / Sdia-r primer pair that bind within the *STA1* ORF (Brandl, 2006), and our newly designed primer pair that bind within the *STA1* promoter. As expected, amplification using the Sdia-f / Sdia-r primer pair was observed for both WY3711 and WLP570 (threshold cycles around 17-19), while no signal above the background was observed for the negative control A15 after 40 cycles (Figure 4B). With the new primer pair on the other hand, amplification was only observed for the highly diastatic WY3711, while no signal above the background was observed for the poorly diastatic WLP570 and the negative control A15 after 40 cycles. We also tested the DNA extracted from the lager yeast cultures contaminated with varying ratios of *S. cerevisiae* WY3711 (10^-6^ – 10^-1^) using the same quantitative PCR reactions. Both primer pairs produced similar linear responses in regard to increasing threshold cycle (C_t_) values as the ratio of WY3711 in the yeast culture decreased 10-fold (Figure 4C). No signal above the background was observed from the yeast culture contaminated with 10^-6^ of *S. cerevisiae* WY3711 after 40 cycles with either of the primer pairs, but the presence of diastatic *S. cerevisiae* could be detected at a concentration of 10^-5^.

### Prevalence of STA1

To determine how common the *STA1* gene is among wild and domesticated strains of *S. cerevisiae,* a BLAST search was performed in the genome assemblies produced in recent whole genome sequencing studies (Gallone et al., 2016; Peter et al., 2018). Because of the chimeric nature of *STA1* (rearranged fragments from *FLO11* and *SGA1*), genomes assembled from short reads have difficulty capturing the full *STA1* sequence on a single contig. To demonstrate, the short-read genome assemblies of three *STA1+* strains (WLP570, OS899 and A81062) were obtained (Krogerus et al., 2016; Peter et al., 2018), and these were searched for the *STA1* sequence (GenBank X02649.1) using BLAST. None of the three assemblies contained the full *STA1* sequence on a single contig (Supplementary Table S3). In addition to short sequence reads, long sequence reads (Nanopore reads for WLP570 and OS899, and PacBio reads for A81062) are available for all three strains (Istace et al., 2017; Krogerus et al., 2016). These were obtained and *de novo* assembled using the LRSDAY pipeline (Yue and Liti, 2018). The long-read genome assemblies were again queried for the *STA1* sequence using BLAST, and now the full sequence was captured on single contigs in all three strains (Supplementary Table S3).

In light of this, *STA1* detection in the short-read genome assemblies was instead carried out by searching for the presence of a 79 bp *STA1*-specific sequence (‘STA1_BLAST’ in Supplementary Table S2). This sequence is amplified by the *STA1*-specific Sdia-f and Sdia-r PCR primers described for the detection of diastatic *S. cerevisiae* (Brandl, 2006). Out of the 1169 publically available genome assemblies that were queried, 54 contained a 100% match to the full 79 bp sequence (Supplementary Table S4). Interestingly, of these 54 strains, 51 were concentrated in the ‘Beer 2’ (‘Mosaic Beer’) and the ‘French Guiana human’ populations, while the remaining three were described as mosaic (Figure 5). While the majority of the strains in these two populations were *STA1+*, not all of them were (69% and 63% in the ‘Beer 2’ and ‘French Guiana human’ populations, respectively).

**Figure 5.**
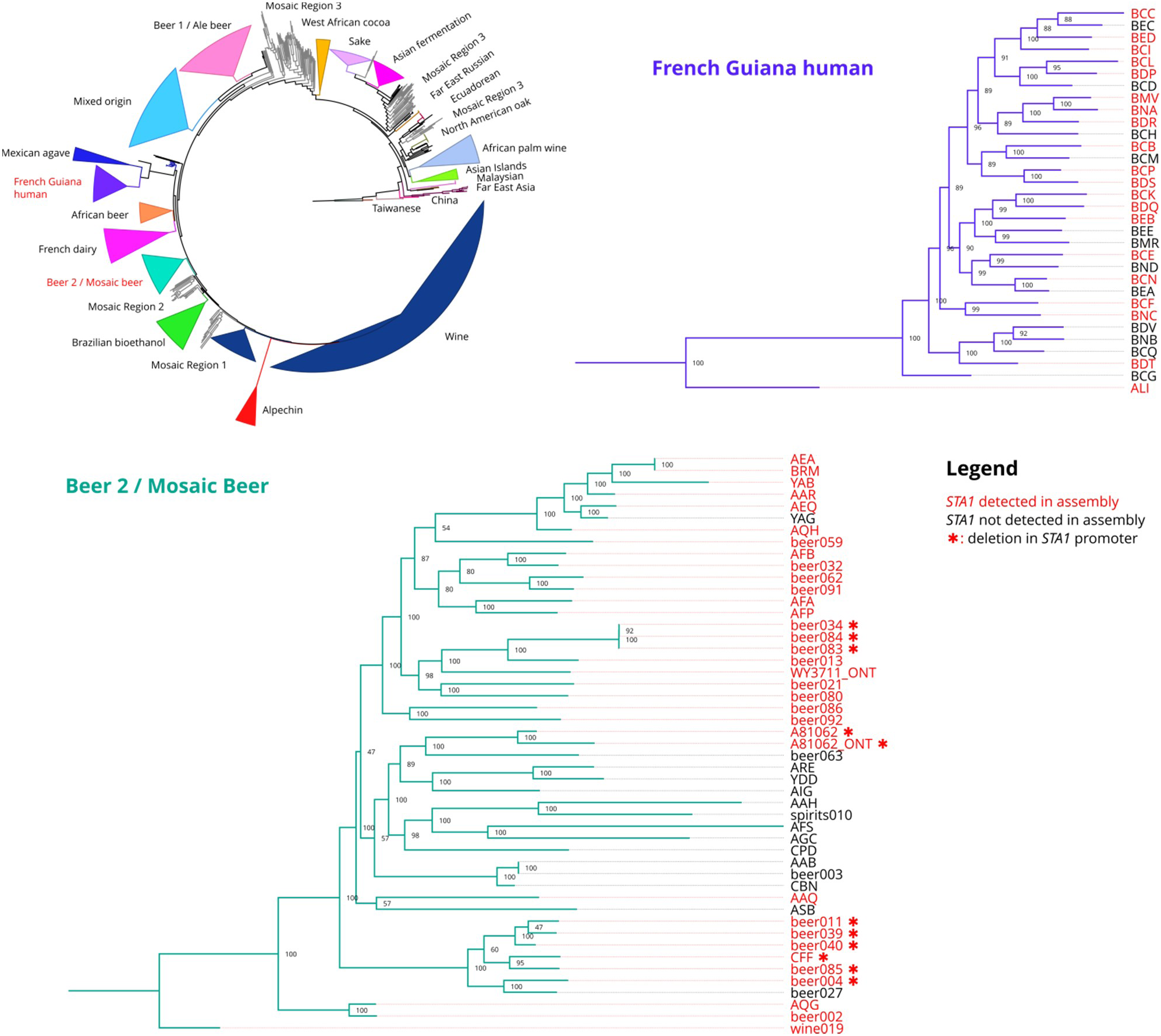
The prevalence of *STA1* in 1171 *S. cerevisiae* genome assemblies. A maximum likelihood phylogenetic tree based on 26 725 238 SNPs at 462842 sites in *S. cerevisiae* genome assemblies obtained from Gallone et al. (2016), Peter et al. (2018) and Krogerus et al. (2016) (rooted with Taiwanese strains as outgroup). Clades have been collapsed to improve clarity and the names of clades containing *STA1+* strains are coloured red (‘French Guiana human’, and ‘Beer 2’/’Mosaic beer’). The ‘French Guiana human’ and ‘Beer 2’/’Mosaic beer’ clades have been expanded, and strain names have been coloured red if *STA1* was detected in the assembly. A red asterisk (*) depicts a strain with a 1162 bp deletion in the *STA1* promoter. The ‘*_ONT’ assemblies were generated in this study from the sequencing reads generated with the Nanopore MinION. Values at nodes indicate bootstrap support values. Branch lengths represent the number of substitutions per site.

The 54 *STA1*+ genomes were also queried for the presence of the 1162 bp deletion in the *STA1* promoter using BLAST (‘STA1_deletion_BLAST’ in Supplementary Table S2). While we observed that the majority of the 15 *STA1+* strains screened in this study had a deletion in the *STA1* promoter, only ten out of the 54 sequenced *STA1*+ strains appeared to have the 1162 bp deletion in the promoter (Figure 5). All ten strains belonged to the ‘Beer 2’ (‘Mosaic Beer’) population.

Whole genome sequencing of *S. cerevisiae* WY3711 also confirmed that the strain belongs to the ‘Beer 2’ (‘Mosaic Beer’) population (Figure 5). The sequencing reads generated with the Nanopore MinION are known to be error-prone (Istace et al., 2017), which naturally affects the reliability of the analysis. Therefore we also sequenced *S. cerevisiae* A62 in the same sequencing run to include as a control. The assembly of A62 using only reads generated from the MinION (polishing with NanoPolish, but no polishing with Illumina reads) showed 99.6% identity with the assembly produced from PacBio reads (polished with Illumina reads), and the assemblies grouped next to each other in the phylogenetic tree (Figure 5).

### STA1 improves maltotriose consumption during wort fermentations

Strains in the ‘Beer 2’ population have been shown to utilize maltotriose efficiently, despite carrying frameshift mutations in *AGT1/MAL11* (Gallone et al., 2016). It has been suggested that the ‘Beer 2’ strains instead utilize alternative mechanisms for maltotriose use. The glucoamylase produced from the *STA1* gene does not only hydrolyse malto-oligomers efficiently, by cleaving *α*-1,4 bonds, but is also active on maltotriose (Kleinman et al., 1988). We therefore explored the role of *STA1* in enabling maltotriose use during wort fermentations.

Initially, we performed fermentations by inoculating 15 °Plato all-malt wort with 1 g L^-1^ of WY3711 and WY3711_D1. During the first 24 hours of fermentation, an identical amount of ethanol was produced by both strains (Figure 6). However, after that time-point, the fermentation with WY3711_D1 slowed considerably. Analysis of the wort sugars during fermentation revealed that maltotriose use in particular, was markedly decreased in WY3711_D1 compared to WY3711 (Figure 6). At 96 hours, for example, WY3711 had consumed nearly 80% of the wort maltotriose, while WY3711_D1 had only consumed 12%. These results suggest that *STA1* has a central role in enabling maltotriose use in *STA1*+ yeast strains.

**Figure 6.**
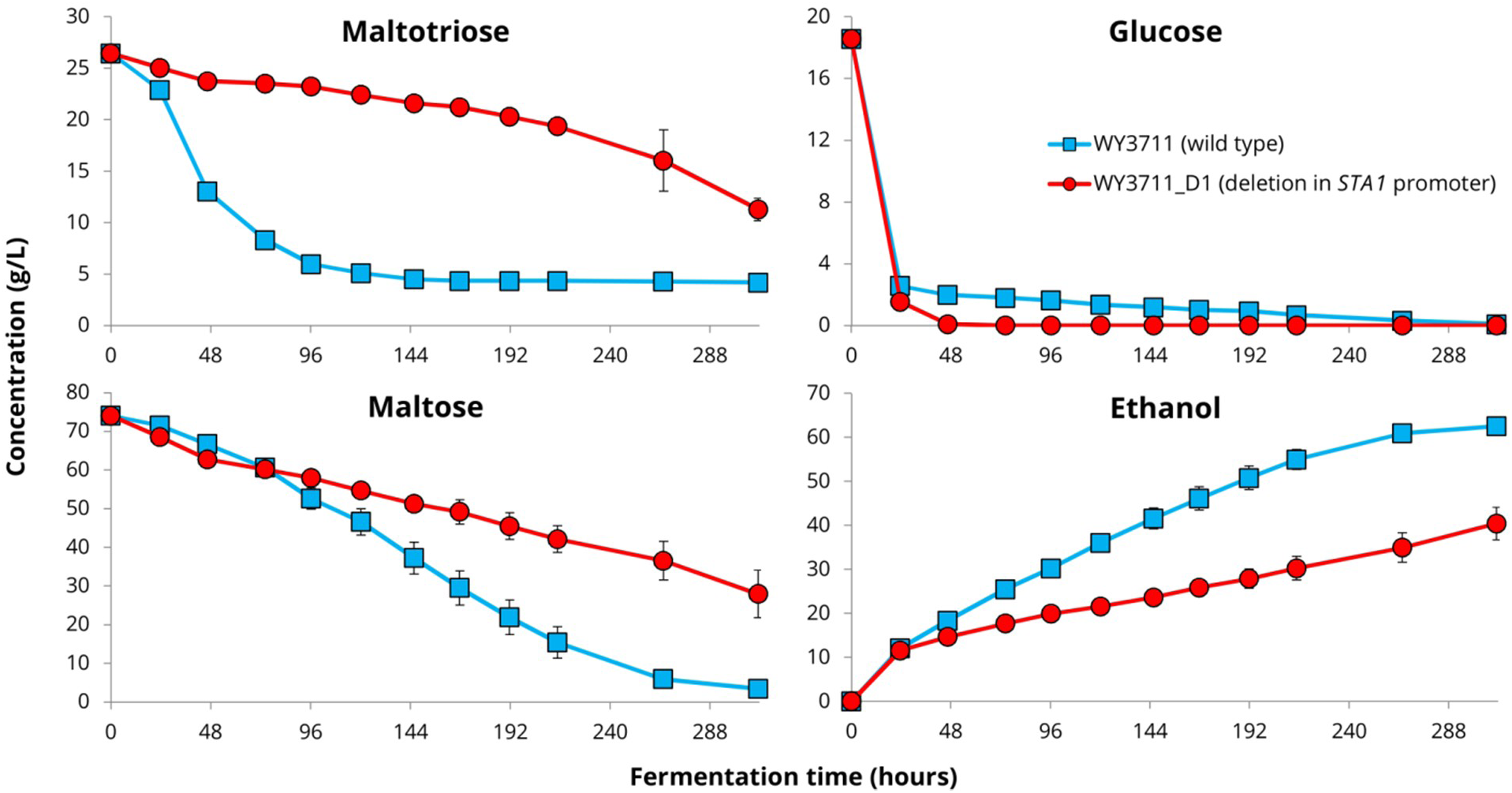
Decreased wort fermentation after the 1162 bp deletion in the *STA1* promoter. The concentrations of fermentable wort sugars and ethanol during fermentations of *S. cerevisiae* WY3711 (blue squares) and WY3711_D1 (1162 bp deletion in *STA1* promoter; red circles) in 15 °P wort. Error bars where visible depict the standard deviation of three replicates.

To ensure that the growth of the WY3711_D1 deletion strain was not impaired from the transformation process, we also compared the growth of WY3711 and WY3711_D1 on YP-Glucose and YP-Maltose in microplate cultivations and saw no significant differences in growth (Supplementary Figure S1). Interestingly, despite showing differential maltotriose use in wort fermentations, we also didn’t observe any significant differences in growth when WY3711 and WY3711_D1 were grown on YNB-Maltotriose in microplate cultivations (Supplementary Figure S1). This would suggest that *STA1* is not the only mechanism enabling maltotriose use in WY3711.

Next we deleted the *STA1* open reading frame from three of the highly diastatic strains: TUM BI PA 109, TUM 71 and WY3711. The *sta1Δ* deletion strains (*_S1 in Table 1) were compared to the wild-type strains in 15 °P wort fermentations. As during the comparative fermentations of WY3711 and WY3711_D1, we observed significantly slower fermentation with the *sta1Δ*, deletion strains compared to the wild-type strains after 24 hours of fermentation (Figure 7A-C). The fermentations with the deletion strains lacking *STA1* also appeared to finish at a lower degree of fermentation than the wild-type strains. Maltotriose use during fermentations was significantly decreased (*p <* 0.001 at the end-point for all strains as determined by two-tailed Student’s t-test) in the strains lacking *STA1* compared to the wild-types (Figure 7D-F). The wild-types of TUM PI BA 109 and TUM 71, for example, had consumed over 80% of the available maltotriose when fermentations were ended, while the *sta1Δ* deletion strains had consumed around 5%. Therefore, it appears as if *STA1* enables efficient consumption of maltotriose from wort and is the main mechanism for maltotriose consumption in these three *STA1+* strains. Interestingly, maltose use was also impaired in both WY3711_D1 (Figure 6) and WY3711_S1 (Figure 7F), suggesting *STA1* also facilitates maltose use in some *STA1+* strains. Nevertheless, the *sta1Δ*, deletion strains still consumed minor amounts of maltotriose, confirming that *STA1* is not the sole mechanism for maltotriose use in these strains. To confirm this we measured the zero-trans uptake rate of maltotriose in these strains using [U-^14^C]-maltotriose. The presence of extracellular glucoamylases should not affect the zero-trans uptake rate, as yeast cells are washed twice prior to the 1-minute exposure to the radio-labelled sugar. Maltotriose uptake ability was detected in all three strains (Figure 7G). As expected, the deletion of *STA1* did not affect the zero-trans maltotriose uptake rate.

**Figure 7.**
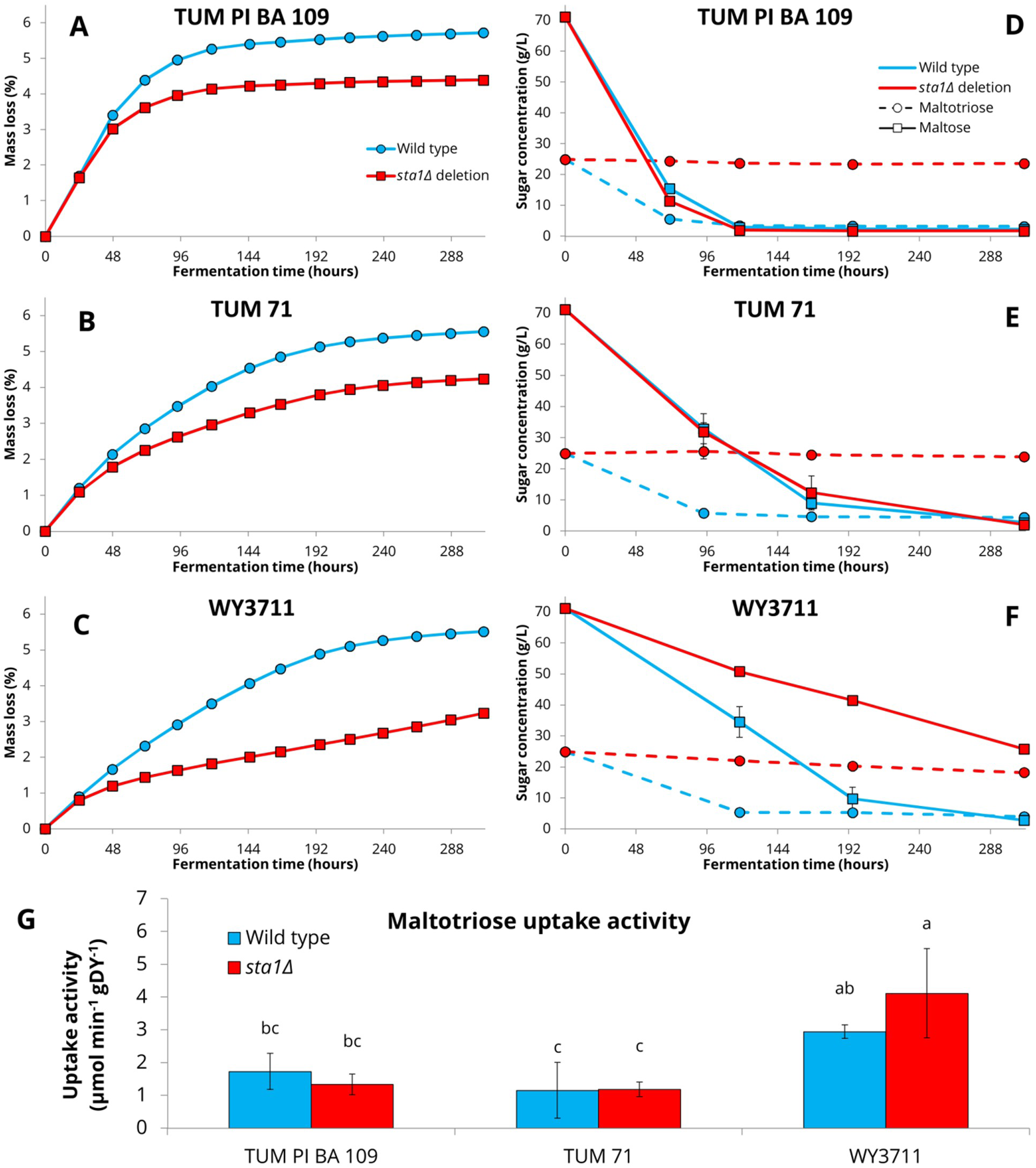
Decreased wort fermentation and maltotriose consumption after deletion of the *STA1* open reading frame in three *STA1+* strains. (**A-C**) The amount of mass lost as CO_2_ (%) during fermentation**. (D-F)** The concentrations (g/L) of maltose (solid line, squares) and maltotriose (dashed line, circles) in the wort during fermentation. (**G**) The zero-trans maltotriose uptake ability (µmol min^-1^ (g dry yeast) ^-1^) of the strains. Values with different letters (a-c) above the bars different significantly (*p* < 0.05 by one-way ANOVA and Tukey’s HSD test) (**A-G**) Wild-type strains in blue, *sta1Δ* deletion strains in red. Error bars where visible depict the standard deviation of three replicates.

We queried the genome assembly of *S. cerevisiae* WY3711 for common maltose and maltotriose transporters (*AGT1*/*MAL11*, *MAL31* and *MTT1*). *MTT1*, a maltotriose transporter with >90% sequence similarity to *MAL31* (Dietvorst et al., 2005), appeared to be present in four copies (>95% sequence identity; Supplementary Table S6) and likely explains the maltotriose uptake ability in WY3711. Despite this, the *sta1Δ* deletion strain WY3711_S1 consumed maltotriose relatively poorly during wort fermentation. The other known maltotriose transporter, *AGT1*/*MAL11,* appeared to be lacking from WY3711 completely, as we were unable to detect it in the genome assembly of WY3711 (Supplementary Table S6) nor did any sequencing reads align to *MAL11* or *MAL13* in the S288C reference genome (Supplementary Figure S3).

We also compared the maltotriose use ability of the nineteen *STA1+* strains (all belonging to the ‘Beer 2’ population) described in Gallone et al. (2016; data was obtained from Supplementary Table S5). We observed that the strains carrying the 1162 bp deletion in the *STA1* promoter, and therefore presumably also exhibiting lower *STA1* expression, had a significantly lower ability to use maltotriose (Supplementary Figure S2; *p* = 0.045, Mann-Whitney U test). This suggests that *STA1* has a central role in enabling maltotriose use in strains of the ‘Beer 2’ (‘Mosaic Beer’) population. However, some strains (e.g. BE034, BE083 and BE084) showed good maltotriose use despite having the deletion in the *STA1* promoter, suggesting that other mechanisms for maltotriose use exist in these strains. This was also confirmed by the fact that all three of the *STA1+* strains that were tested in this study, showed zero-trans uptake rates for maltotriose.

## Discussion

In this study we aimed to elucidate genetic determinants behind the variable diastatic ability that has been observed in *S. cerevisiae* strains carrying the *STA1* gene (i.e. strains previously known as *Saccharomyces diastaticus*). While no nonsense or frameshift mutations were observed in the *STA1* open reading frames of strains that were screened here, we show that multiple *STA1+* strains have a 1162 bp deletion in the *STA1* promoter. The strains with the deletion showed significantly less growth in beer and on starch, and consumed less dextrin when grown in media with dextrin as the sole carbon source. Reverse engineering of the most active strain confirmed the role of this deletion in decreasing the diastatic ability.

Diastatic *S. cerevisiae* remains a widespread and important contaminant of beer, particularly in smaller breweries where beers are seldom pasteurized, quality control is less stringent, and experimentation with different yeast strains is more common (Meier-Dörnberg et al., 2018). Using the results of this study, we developed new PCR primers which can differentiate between highly active spoilage strains harbouring the full *STA1* promoter, and less active strains with the deletion in the promoter. This improves the reliability of the detection methods and can potentially reduce waste and unnecessary costs related to product recalls caused by false positives. Differentiation between highly active and benign diastatic *S. cerevisiae* has currently only been possible with time-consuming plate-based microbiological methods (van der Aa Kühle, 1998).

The variable diastatic ability in *STA1+* strains appears to be a result of variable gene expression. Genomic studies have shown that mutations in non-coding regions play a large role in determining phenotype diversity by affecting gene regulation (Almeida et al., 2017; Connelly et al., 2013). Here, the deleted region contains an upstream activation sequence and transcription factor (Ste12 and Tec1) binding site (Kim et al., 2004a, 2004b). This upstream activation sequence has been shown to facilitate gene expression (Kim et al., 2004a, 2004b). We saw over 100-fold higher expression of *STA1* in a strain containing the full promoter, compared to two strains lacking the 1162 bp region in the promoter (natural and CRISPR/Cas9-mediated deletion), suggesting that regulation of *STA1* is the cause of the differences in diastatic ability that were observed between the strains with the full and partial *STA1* promoter. The results observed here highlight the importance of intergenic polymorphisms and their potential effect on phenotype. Such mutations are commonly overlooked, for example in adaptive evolution studies (Krogerus et al., 2018; Quarterman et al., 2016; Wallace-Salinas et al., 2015), and warrant more emphasis in future studies.

Wort fermentations with the engineered strains revealed that the *STA1*-encoded glucoamylase appears to have a central role in enabling the use of oligomeric wort sugars, such as maltotriose, by *STA1*+ strains during wort fermentation. It has previously been assumed that transport of maltotriose into the yeast cells is required for it to be consumed during wort fermentations (Alves et al., 2008; Day et al., 2002; Rautio and Londesborough, 2003), but our results suggest that extracellular hydrolysis of maltotriose through *STA1* allows for efficient consumption of maltotriose from the wort throughout fermentation. A recent study also reported extracellular hydrolysis of maltotriose by an *α*-glucosidase encoded by *IMA5* (Alves et al., 2018), however, in that case maltotriose hydrolysis was only observed after an extensive lag phase. In brewing yeast where maltotriose use is enabled through the *AGT1*-encoded transporter, maltotriose is typically consumed from the wort only towards the later parts of fermentation when maltose concentrations are low (Rautio and Londesborough, 2003). Here, in the *STA1+* strains, maltotriose was consumed rapidly from the wort alongside maltose.

Brewers have traditionally associated diastatic contaminants with ‘wild’ yeast (Meier-Dörnberg et al., 2018), but here we show that this trait is associated with two domesticated *S. cerevisiae* populations, ‘Beer 2’/’Mosaic beer’ and ‘French Guiana human’ (Gallone et al., 2016; Peter et al., 2018). Strains of both the ‘Beer 1’/’Ale beer’ and ‘Beer 2’/’Mosaic beer’ populations have been shown to utilize maltotriose efficiently, and this trait is considered a domestication signature of brewing strains (Gallone et al., 2016). Maltotriose use in the ‘Beer 1’ strains has been linked to the presence and increased copy number of *AGT1/MAL11,* but this allele is either absent or non-functional in the ‘Beer 2’/’Mosaic beer’ strains (Gallone et al., 2016). Our results suggest that *STA1* is the, until now, unknown mechanism that enables efficient maltotriose use in the ‘Beer 2’/’Mosaic beer’ strains. We propose that the formation and retention of *STA1,* through the chimerization of *FLO11* and *SGA1* (Lo and Dranginis, 1996; Yamashita et al., 1987), is an alternative evolutionary strategy for efficient utilization of sugars present in brewer’s wort. The presence of *STA1* is expected to provide a fitness advantage to strains lacking the ability to use maltotriose in a wort environment, and this should be tested in future studies. While *MTT1* also appears to be present in many ‘Beer 2’/’Mosaic beer’ strains, the results here suggest that extracellular hydrolysis of maltotriose seems to be the dominant route for utilization. It may be speculated that this trait was later counter-selected by brewers, mediated by the deletion in the promoter. This counter-selection may have been driven by the desired specifications of the beer (e.g. not overly dry) or the need to store beer for extended periods without excessive build-up of pressure in vessels. Interestingly, chimerization of two *SeMALT* genes to form a functional maltotriose transporter was recently also been demonstrated in two adaptive evolution studies on maltotriose fermentation (Baker and Hittinger, 2019; Brouwers et al., 2019).

The prevalence of *STA1* in the ‘French Guiana human’ population (Peter et al., 2018) was unexpected. These isolates were obtained from various origins including fruits, animals, but mainly human faeces (Angebault et al., 2013; Peter et al., 2018). The preparation and consumption of cachiri, a traditional starch-rich fermented beverage made from cassava (Barre, 1938; Carrizales et al., 1986), is widespread among the humans that were sampled (Angebault et al., 2013), and it is possible that *STA1* also provides a fitness advantage in the starch-rich environment of cachiri. The possible link between the ‘Beer 2’/’Mosaic beer’ and ‘French Guiana human’ strains remains unclear, and this should be clarified in future studies. The sequence similarity of the *STA1* gene from *S. cerevisiae* WY3711 (‘Beer 2’) and OS899 (‘French Guiana human’) around the *FLO11*/*SGA1* junction and in the promoter suggests that they might have a common origin (Supplementary Figure S6).

In conclusion, we show here that the variable diastatic ability that has been observed in *STA1+* strains is a result of a deletion in the *STA1* promoter, and that *STA1* also plays a central role in enabling maltotriose use during wort fermentations. This allows for the improved reliability of molecular detection methods for diastatic contaminants in beer, and can be exploited for strain development where maltotriose use is desired. To further clarify the role of *STA1* as a mechanism enabling maltotriose use in brewing strains, its effect on fitness in a brewing environment should be elucidated. In addition, the ability of *STA1* to enable maltotriose use could be confirmed by expressing *STA1* in a strain lacking zero-trans maltotriose uptake ability.

## Acknowledgements

We thank Mathias Hutzler and Richard Preiss for providing strains. This work was supported by the Alfred Kordelin Foundation, Svenska Kulturfonden - The Swedish Cultural Foundation in Finland, Suomen Kulttuurirahasto, and the Academy of Finland (Academy Project 276480).

## Author Contributions Statement

KK conceived the study. KK, JK, and BG designed the experiments. KK, FM and JK conducted the experiments described in this study. KK analysed all data. KK wrote the manuscript. BG, FM and JK edited the manuscript. All authors read and approved the final manuscript.

## Data Availability Statement

The datasets generated for this study can be found in the NCBI’s Short Read Archive under BioProject PRJNA544899 in the NCBI BioProject database (https://www.ncbi.nlm.nih.gov/bioproject/).

## Supplementary Figures and Tables

**Figure S1.**
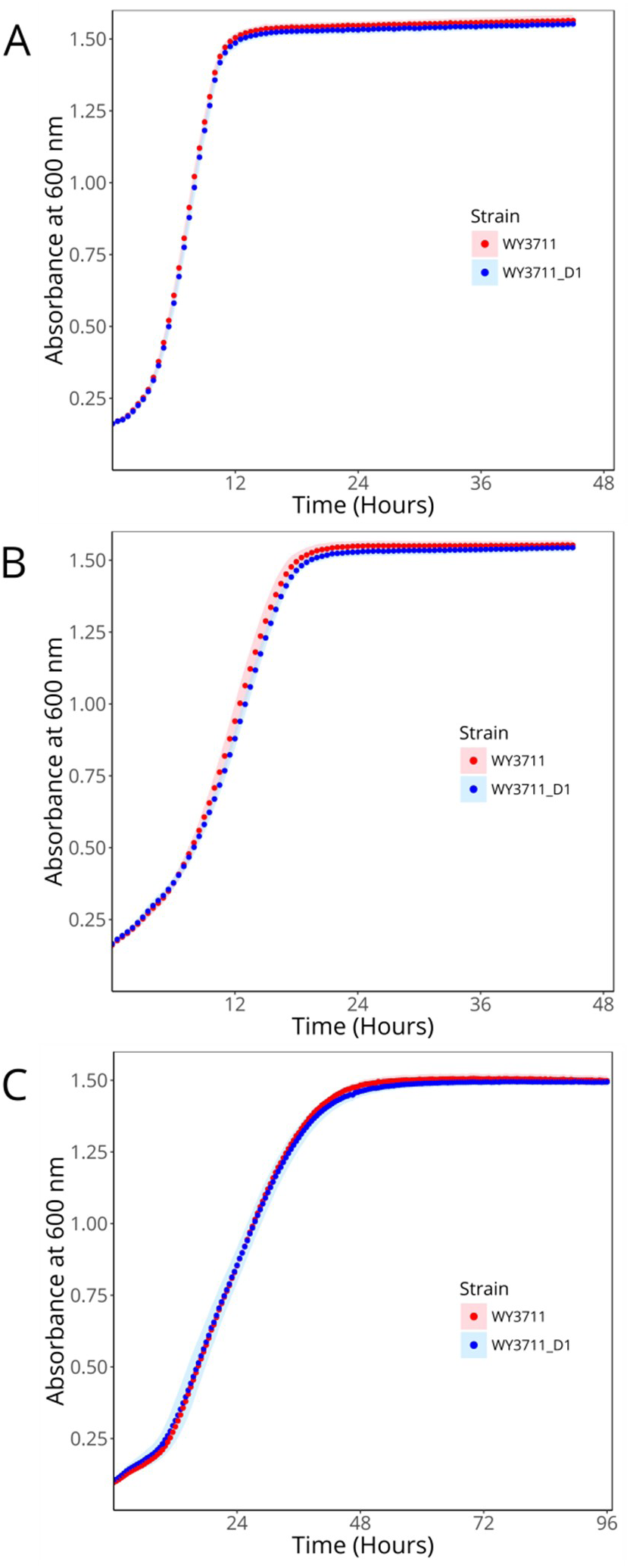
The growth (absorbance at 600 nm) of *S. cerevisiae* WY3711 (red) and WY3711_D1 (blue) in **(A)** YP-Glucose (1%), (**B**) YP-Maltose (1%), and (**C**) YNB-Maltotriose (1%). Cultivations were performed in microplate format at 25 °C. Points and shaded areas represent the mean and standard deviation of 8 biological replicates per strain, respectively. No significant difference was observed between the two strains in any media (two-tailed Student’s t-test, *p* > 0.05).

**Figure S2.**
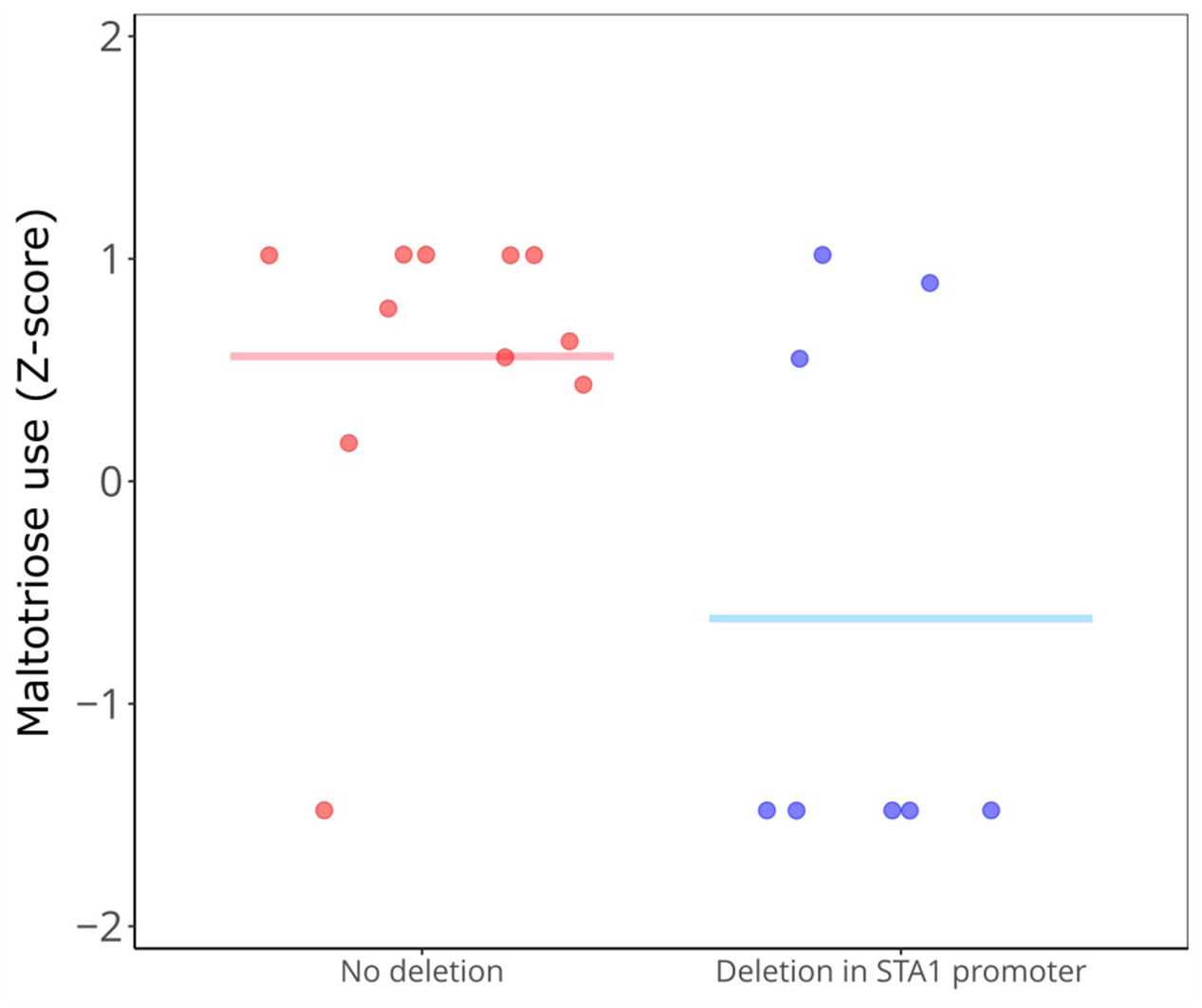
The ability to use maltotriose among the *STA1+* strains studied by Gallone et al. (2016). Strains are grouped depending on whether they have an 1162 bp deletion in the *STA1* promoter. Z-scores were obtained from Supplementary Table S5 in Gallone et al. (2016). The group average is depicted as a straight line. The two groups differed significantly (Mann-Whitney U test, *p* = 0.045).

**Figure S3.**
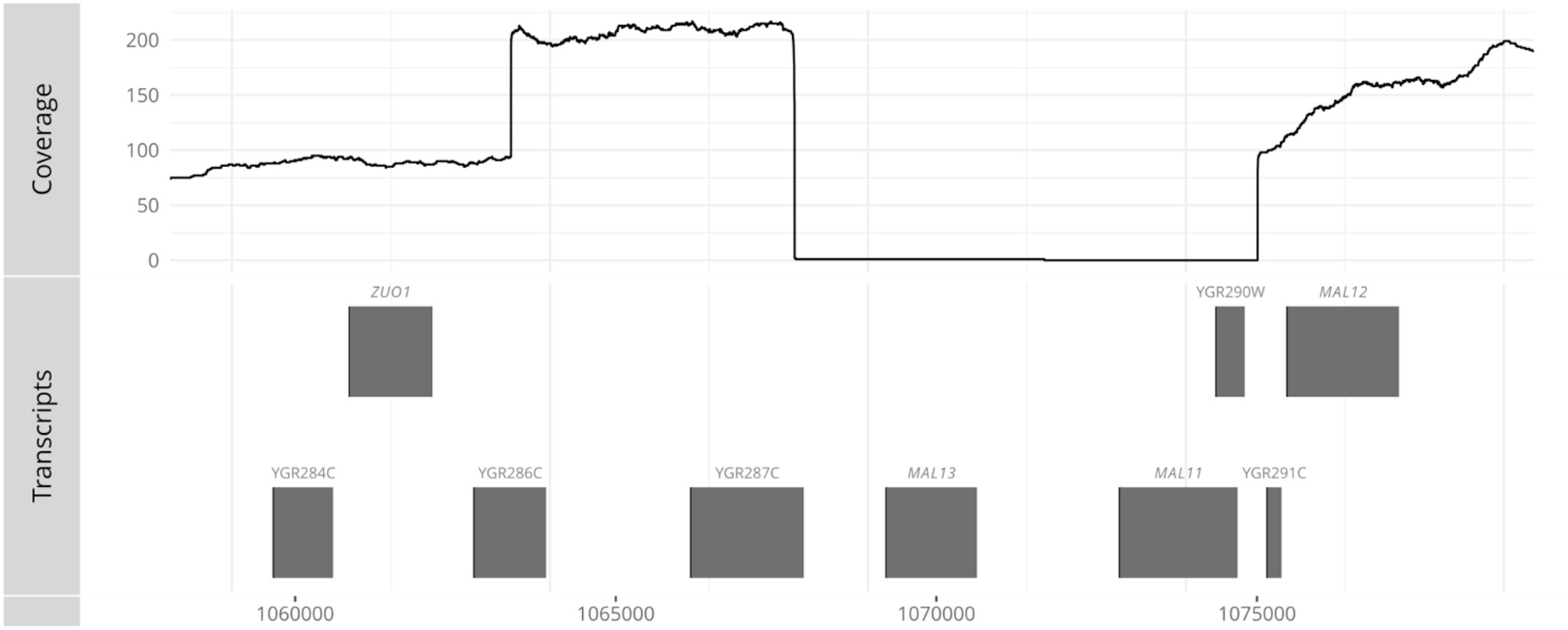
The sequencing coverage of Nanopore reads from *S. cerevisiae* WY3711 aligned to *S. cerevisiae* S288C around the *MAL1* locus (chromosome VII: 1060000-1080000).

**Figure S4.**
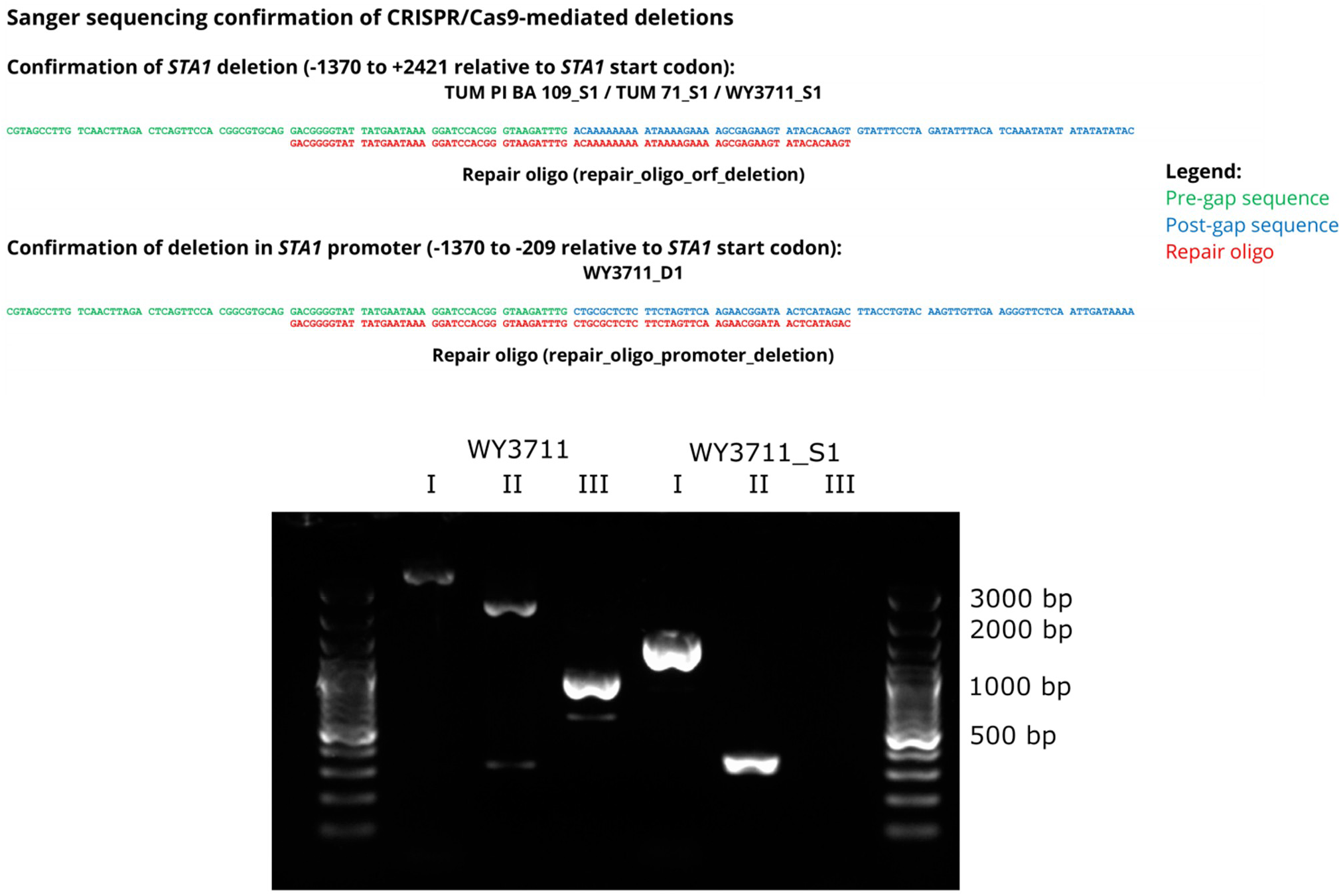
Confirmation of CRISPR/Cas9-mediated deletions by Sanger sequencing and PCR. Primers used to confirm the deletion of *STA1* (−1370 to +2421 relative to start codon) by PCR: **I**: STA1_Full_Fw / STA1_Full_Rv, **II**: STA1_1055_F / STA1_5201_R, **III**: SD-5A / SD-6B

**Figure S5.**
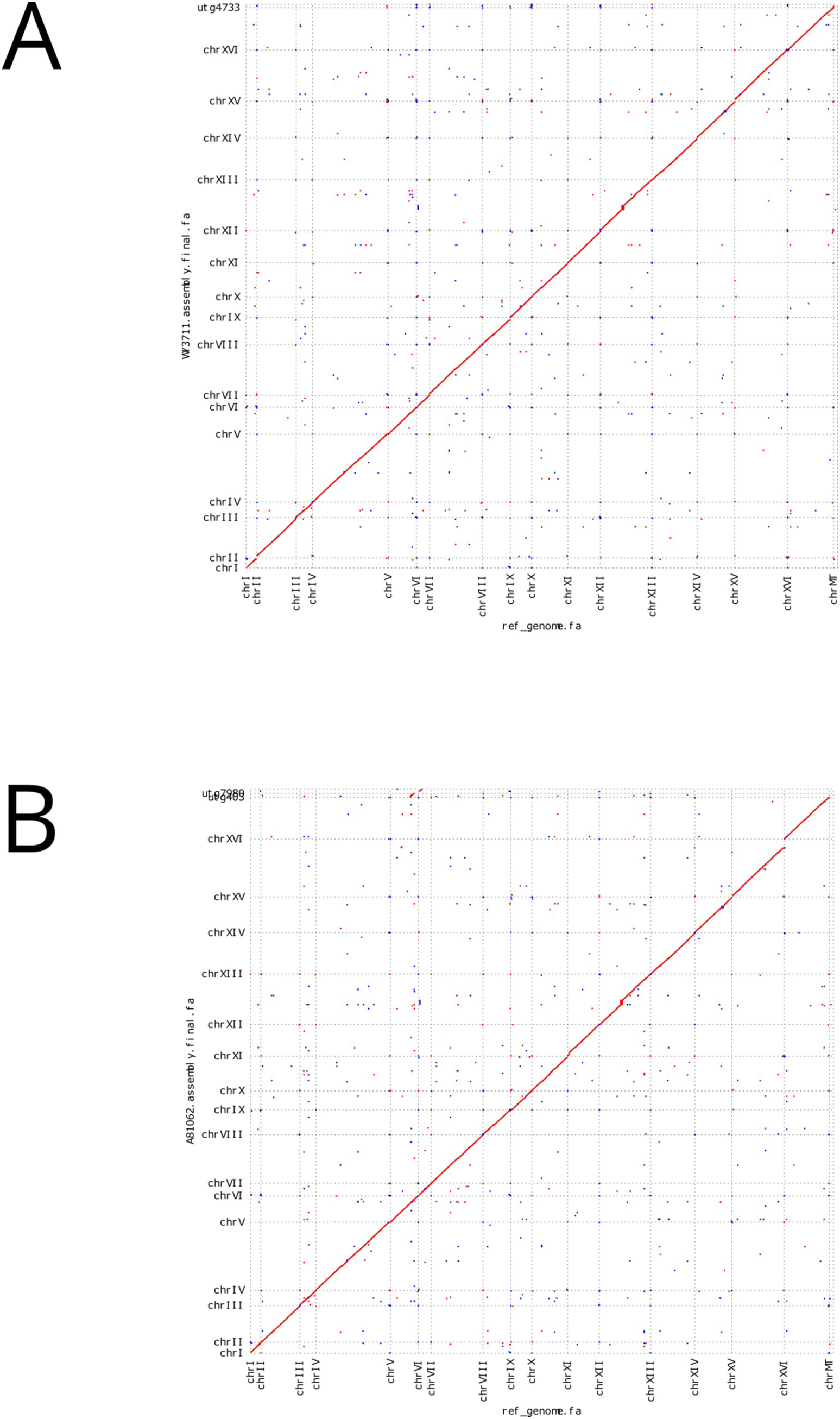
Comparison of (**A**) *S. cerevisiae* WY3711 and (**B**) *S. cerevisiae* A81062 *de novo* assemblies with *S. cerevisiae* S288C reference genome.

**Figure S6.**
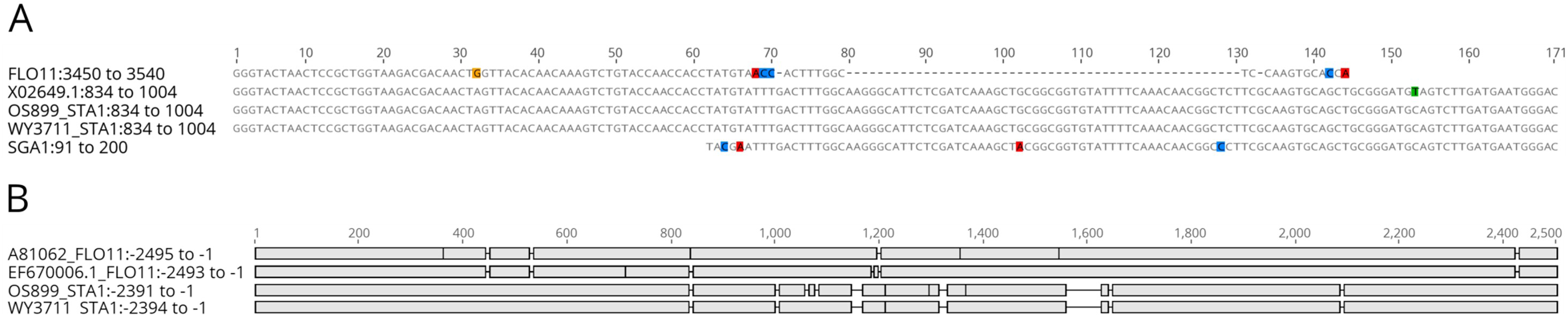
Multiple sequence alignment of (**A**) sequences around the *FLO11*/*SGA1* junction in *STA1* (Genbank X02649.1) from *S. cerevisiae* WY3711 (‘Beer 2’/’Mosaic Beer’) and *S. cerevisiae* OS899 (‘French Guiana human’), and (**B**) sequences upstream of *STA1* from *S. cerevisiae* WY3711 (‘Beer 2’/’Mosaic Beer’) and *S. cerevisiae* OS899 (‘French Guiana human’) and *FLO11* (Genbank EF670006.1) from *S. cerevisiae* A81062 (‘Beer 2’/’Mosaic Beer’).

**Supplementary Table S1.**
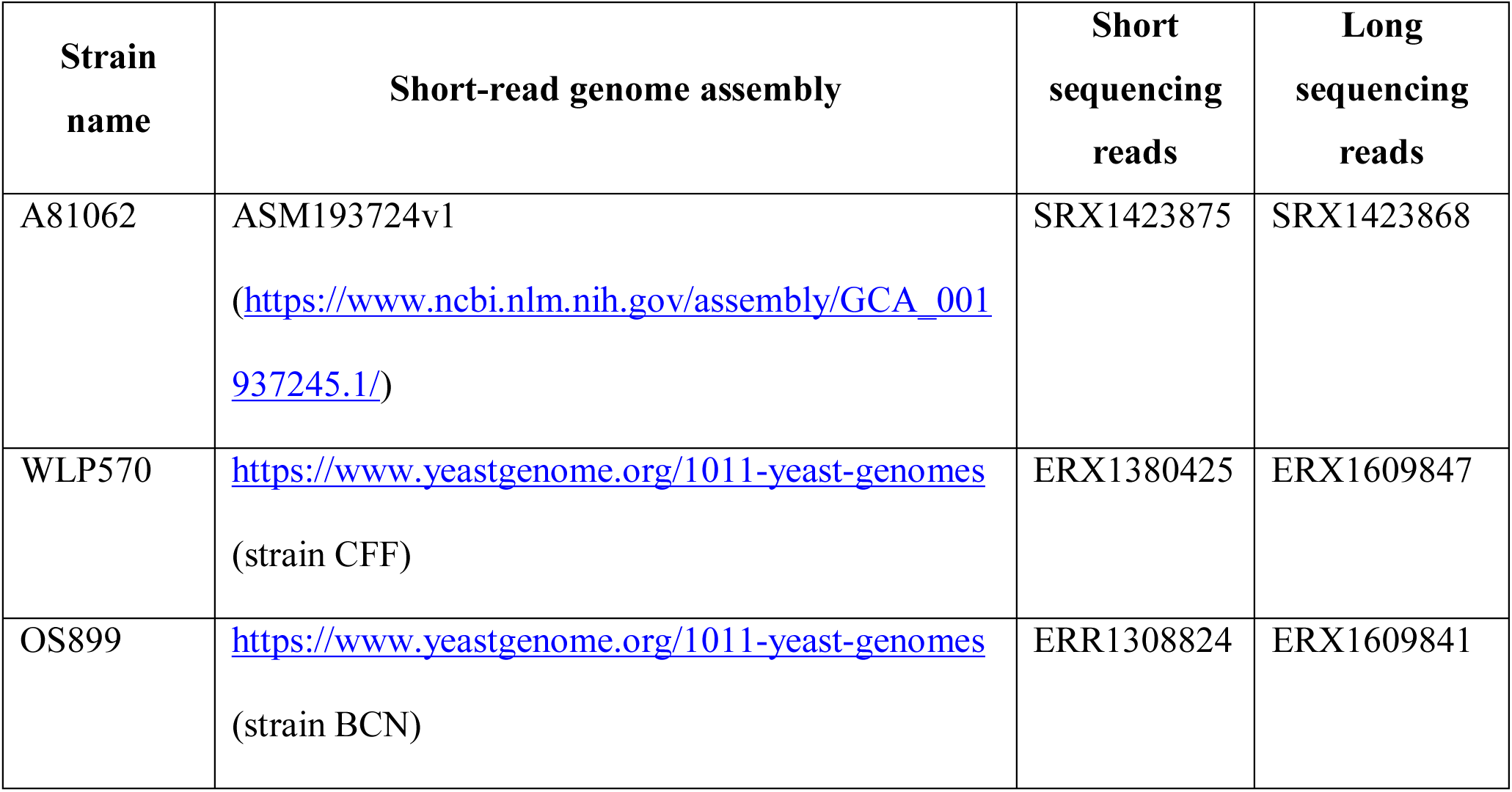
Accession numbers or links to genome assemblies, short sequencing reads, and long sequencing reads of three *STA1+* strains.

**Supplementary Table S2.**
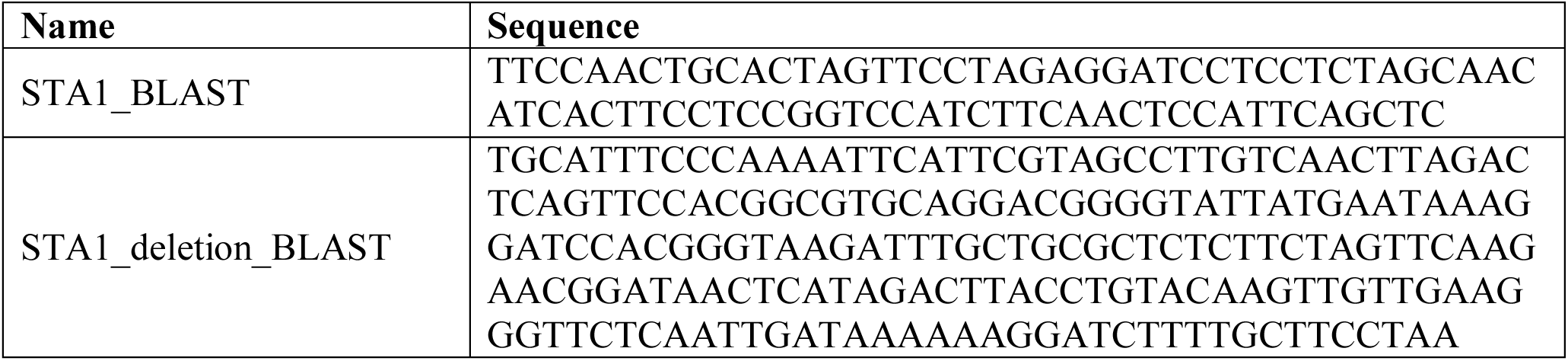
The sequences used to query for the presence of *STA1* and the 1162 bp deletion in the *STA1* promoter.

**Supplementary Table S3.**
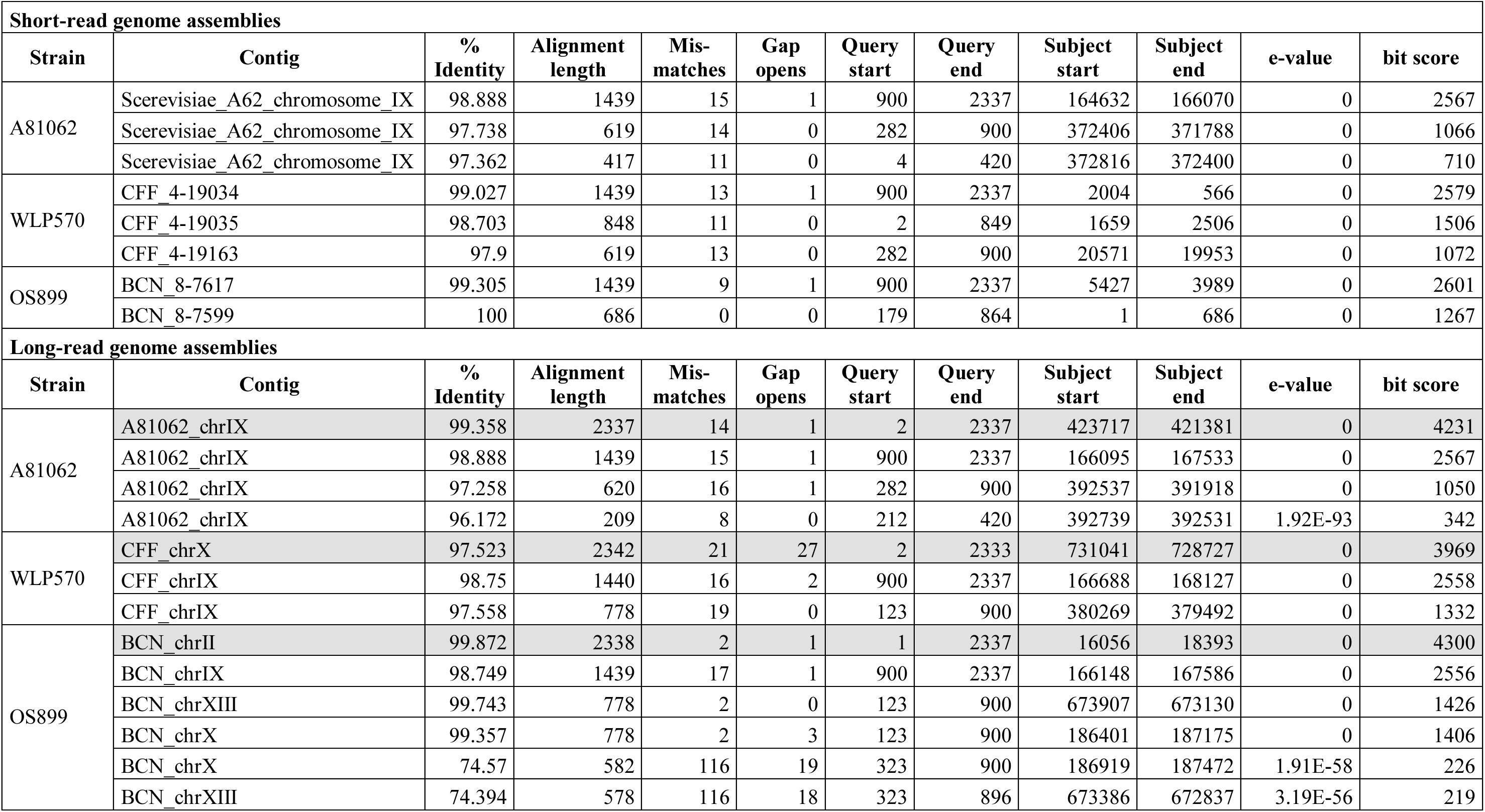
BLAST results for *STA1* (Genbank X02649.1). The full-length hits have been highlighted with a light grey background.

**Supplementary Table S4.**
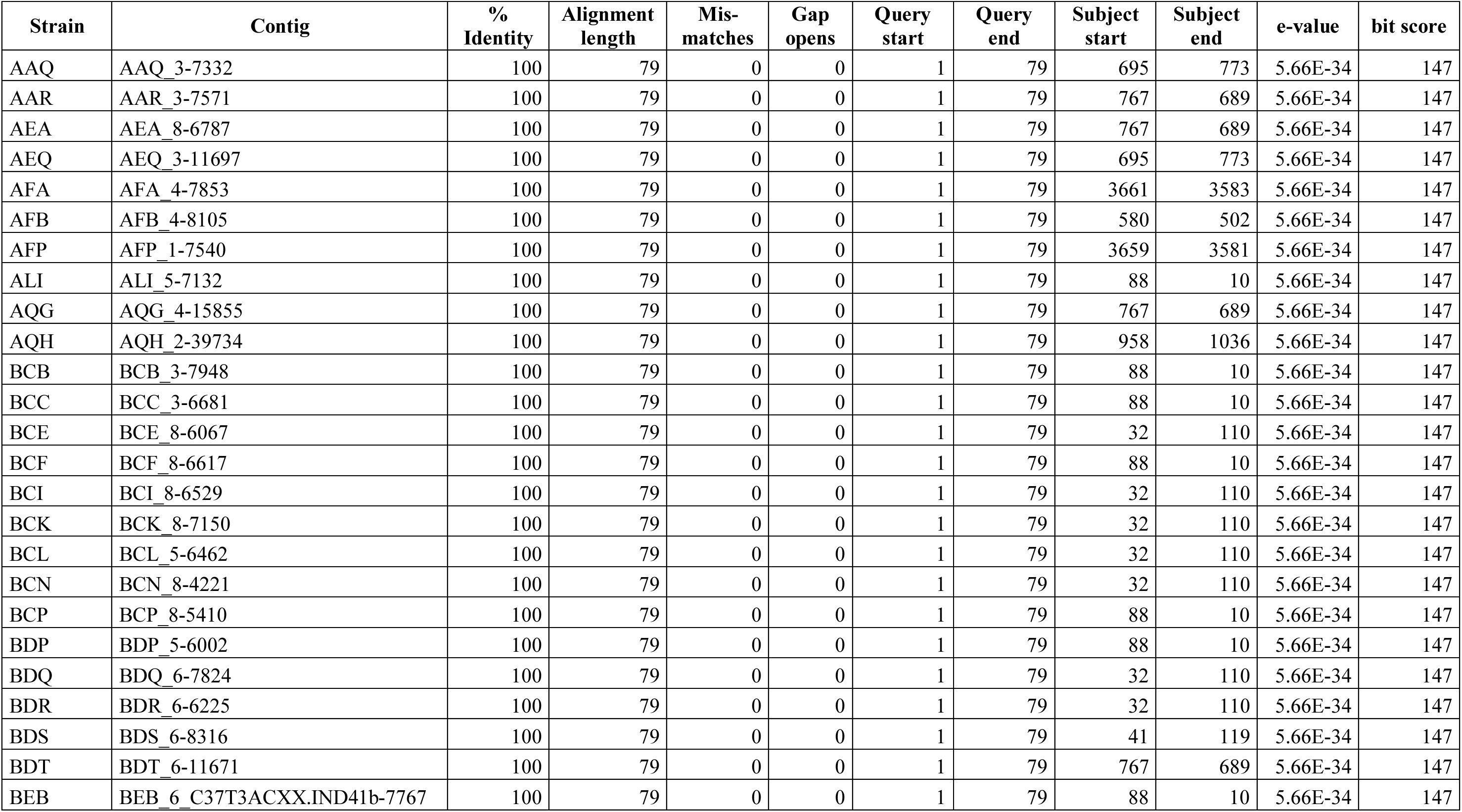
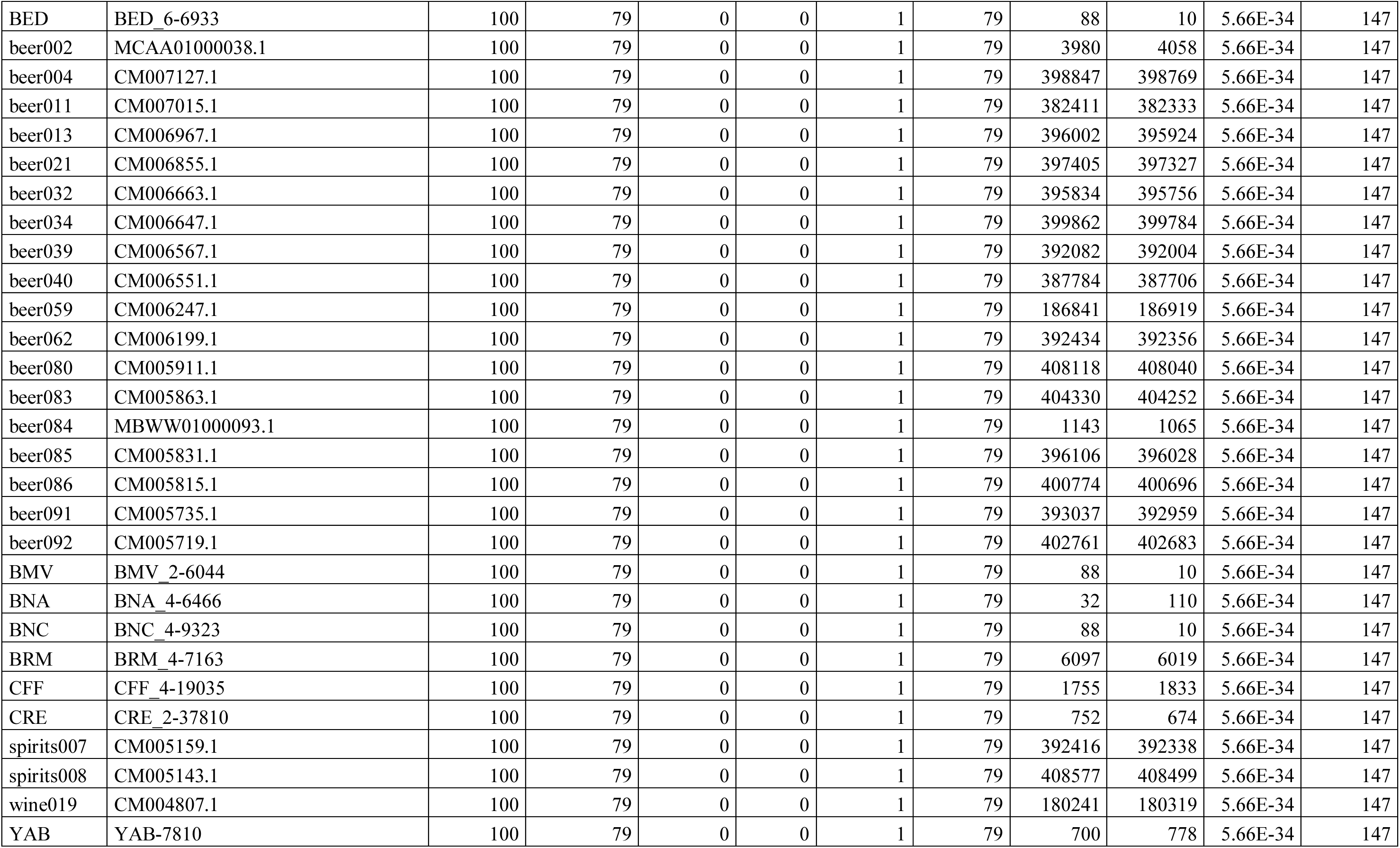
BLAST results for ‘STA1_BLAST’ (Supplementary Table S2) in the 1169 *S. cerevisiae* genome assemblies from Gallone et al. (2016) and Peter et al. (2018).

**Supplementary Table S5.**
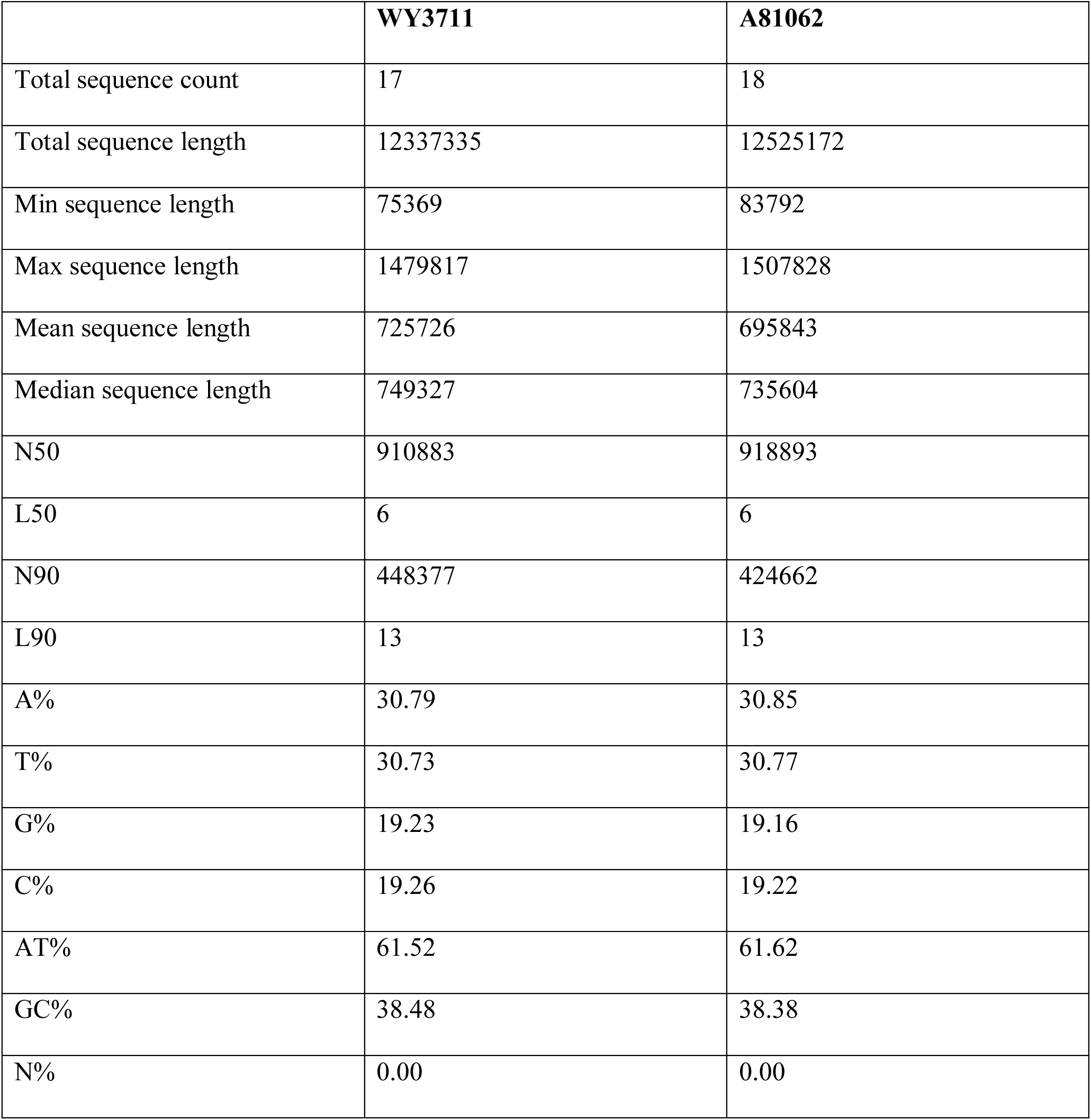
The assembly statistics for *S. cerevisiae* WY3711 and A81062

**Supplementary Table S6.**
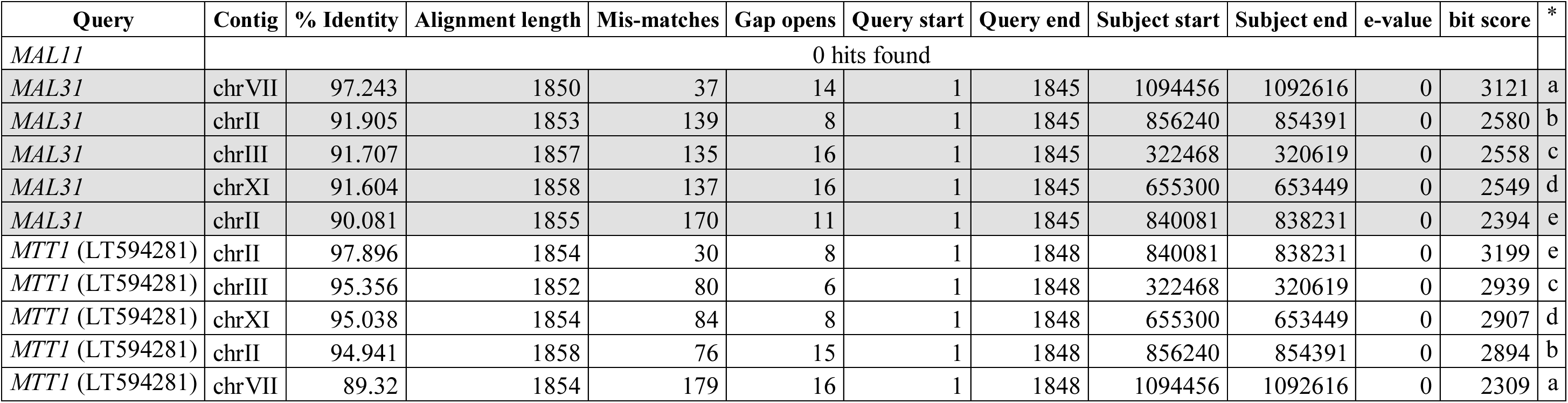
BLAST results for *MAL11*, *MAL31* and *MTT1* (GenBank LT594281) in *S. cerevisiae* WY3711. Hits from different queries with the same letter in final column (*) are the same subject sequence.

