## Supplementary material for "A deletion in the *STA1* promoter determines maltotriose and starch utilization in *STA1*+ *Saccharomyces cerevisiae* strains": readme.txt

Supplementary_Data.fasta contains the sequences of the STA1 open reading frame and upstream region in the 15 Saccharomyces cerevisiae strains that were analysed in the study.
A81062.ONT_only.final.fa is the de novo assembly of S. cerevisiae A81062 generated only from reads produced with the MinION (polished with NanoPolish).
WY3711.ONT_only.final.fa is the de novo assembly of S. cerevisiae WY3711 generated only from reads produced with the MinION (polished with NanoPolish).
