## Supplementary Figures and Tables for "A deletion in the *STA1* promoter determines maltotriose and starch utilization in *STA1*+ *Saccharomyces cerevisiae* strains"

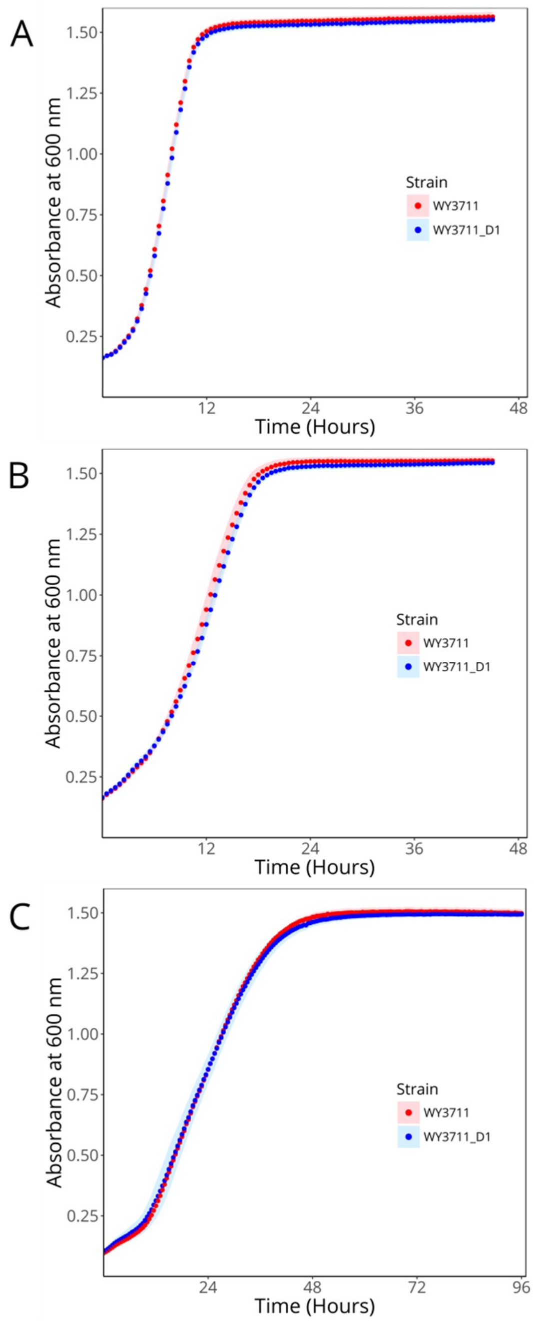


**Figure S1** – The growth (absorbance at 600 nm) of *S. cerevisiae* WY3711 (red) and WY3711_D1 (blue) in **(A)** YP-Glucose (1%), (**B**) YP-Maltose (1%), and (**C**) YNB-Maltotriose (1%). Cultivations were performed in microplate format at 25 °C. Points and shaded areas represent the mean and standard deviation of 8 biological replicates per strain, respectively. No significant difference was observed between the two strains in any media (two-tailed Student's t-test, $p > 0.05$).

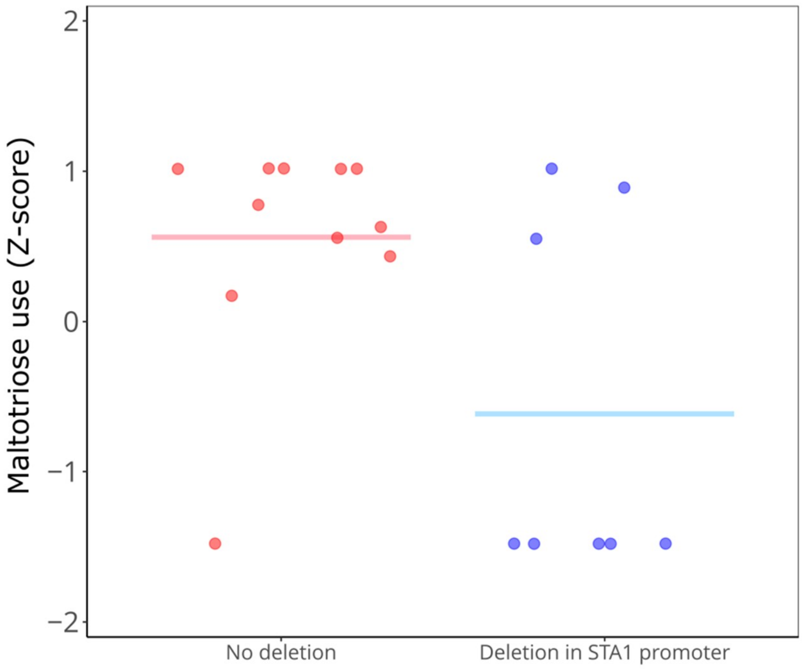


**Figure S2** – The ability to use maltotriose among the *STA1+* strains studied by Gallone et al. (2016). Strains are grouped depending on whether they have an 1162 bp deletion in the *STA1* promoter. Z-scores were obtained from Supplementary Table S5 in Gallone et al. (2016). The group average is depicted as a straight line. The two groups differed significantly (Mann-Whitney U test, $p = 0.045$).

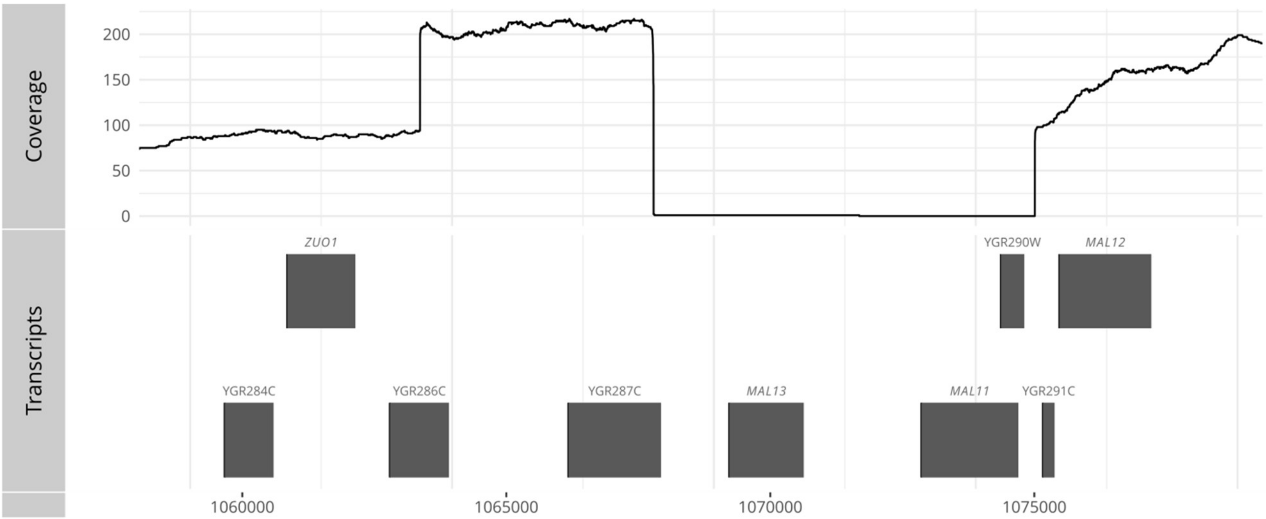


**Figure S3** – The sequencing coverage of Nanopore reads from *S. cerevisiae* WY3711 aligned to *S. cerevisiae* S288C around the *MAL1* locus (chromosome VII: 1060000-1080000).

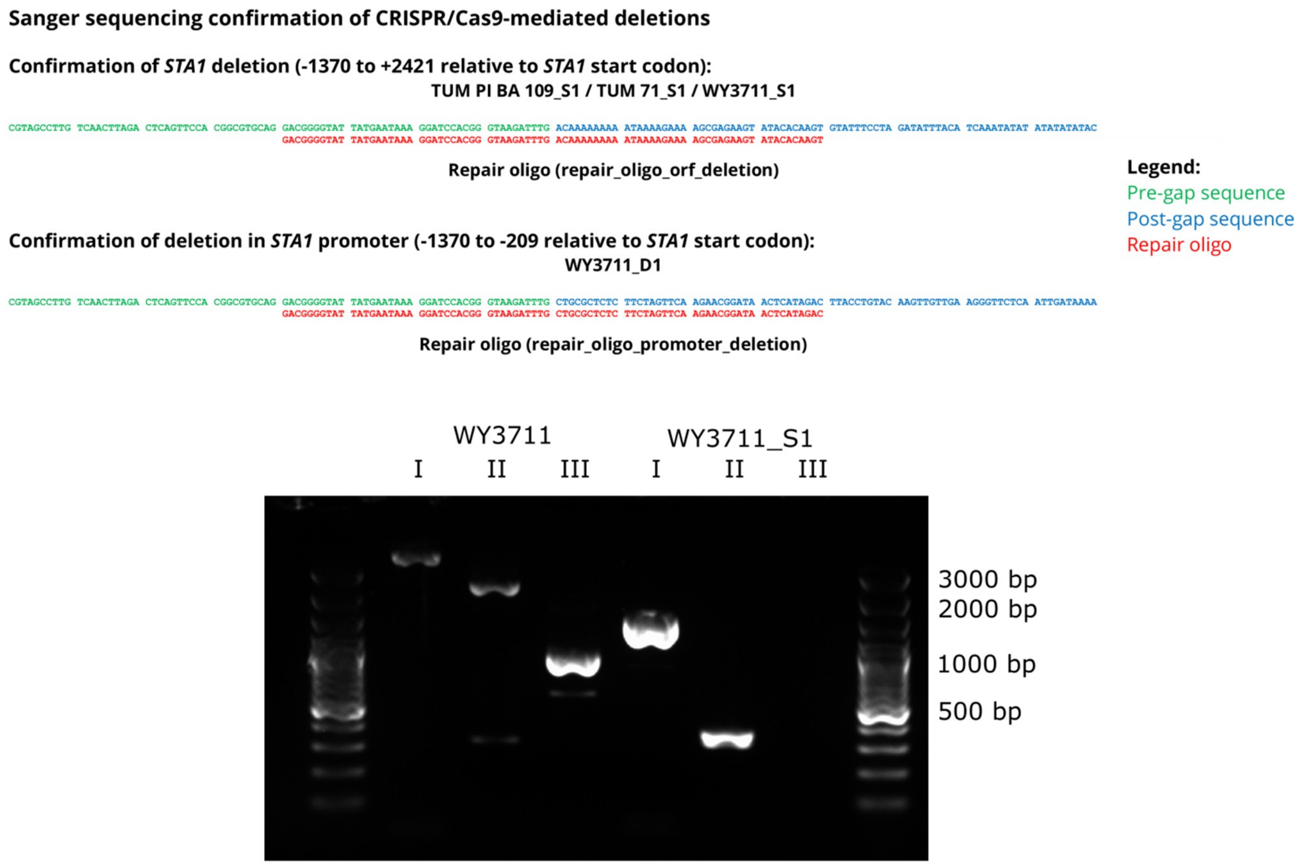


**Figure S4** – Confirmation of CRISPR/Cas9-mediated deletions by Sanger sequencing and PCR. Primers used to confirm the deletion of *STA1* (-1370 to +2421 relative to start codon) by PCR: **I**: STA1_Full_Fw / STA1_Full_Rv, **II**: STA1_1055_F / STA1_5201_R, **III**: SD-5A / SD-6B

A

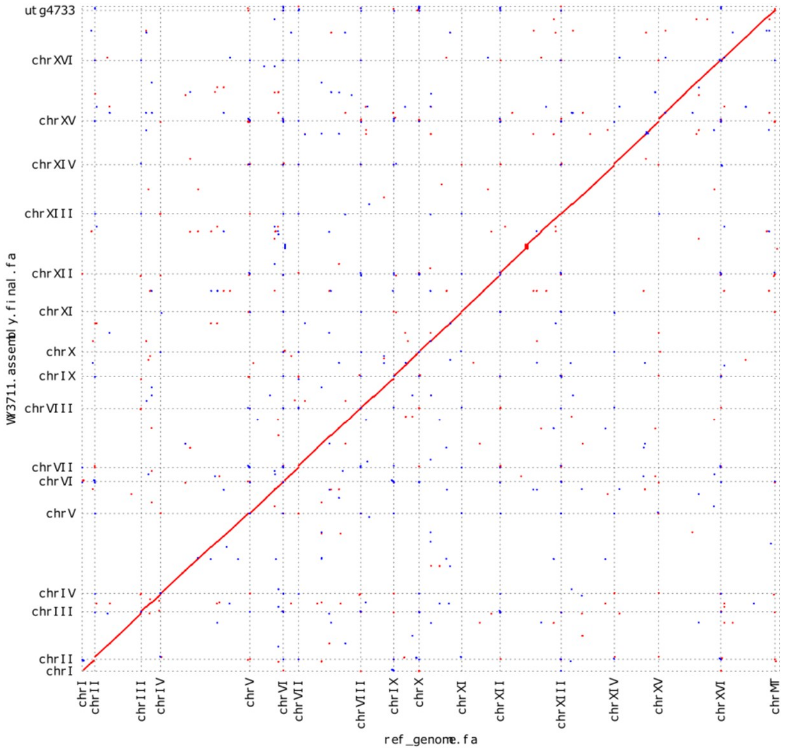


B

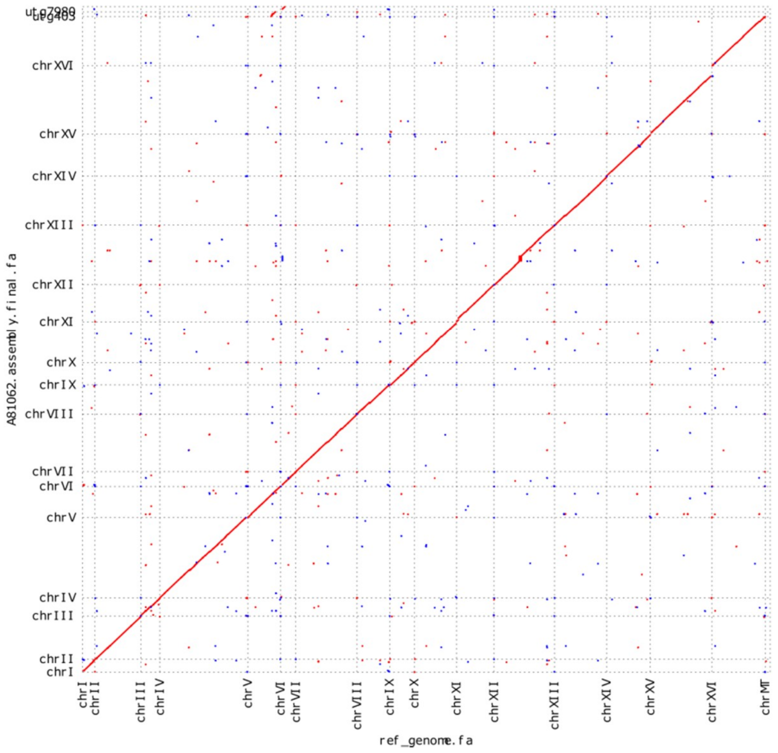


**Figure S5** – Comparison of (**A**) *S. cerevisiae* WY3711 and (**B**) *S. cerevisiae* A81062 *de novo* assemblies with *S. cerevisiae* S288C reference genome.

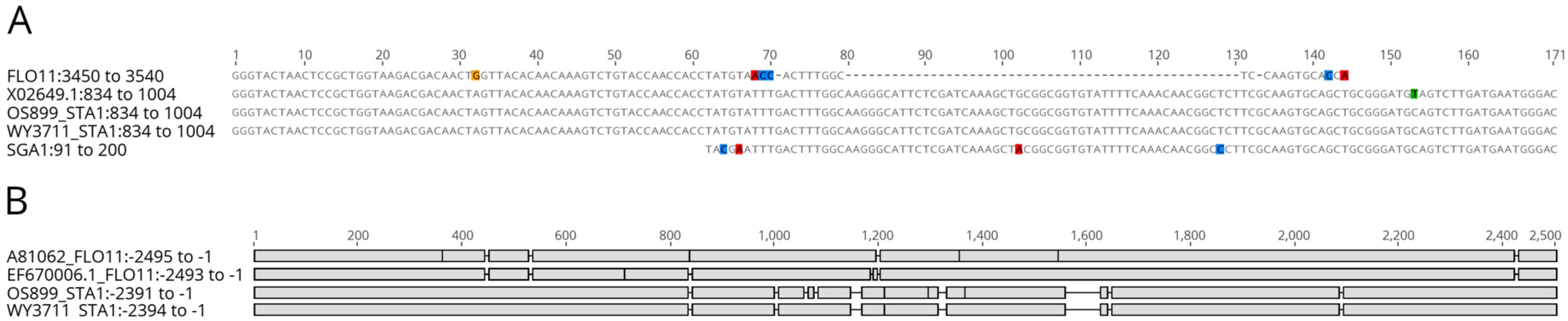


**Figure S6** – Multiple sequence alignment of (**A**) sequences around the *FLO11*/*SGA1* junction in *STA1* (Genbank X02649.1) from *S. cerevisiae* WY3711 (‘Beer 2’/’Mosaic Beer’) and *S. cerevisiae* OS899 (‘French Guiana human’), and (**B**) sequences upstream of *STA1* from *S. cerevisiae* WY3711 (‘Beer 2’/’Mosaic Beer’) and *S. cerevisiae* OS899 (‘French Guiana human’) and *FLO11* (Genbank EF670006.1) from *S. cerevisiae* A81062 (‘Beer 2’/’Mosaic Beer’).

**Supplementary Table S1** – Accession numbers or links to genome assemblies, short sequencing reads, and long sequencing reads of three *STA1+* strains.

| Strain name | Short-read genome assembly | Short sequencing reads | Long sequencing reads |
|---|---|---|---|
| A81062 | ASM193724v1 (https://www.ncbi.nlm.nih.gov/assembly/GCA_001937245.1/) | SRX1423875 | SRX1423868 |
| WLP570 | https://www.yeastgenome.org/1011-yeast-genomes (strain CFF) | ERX1380425 | ERX1609847 |
| OS899 | https://www.yeastgenome.org/1011-yeast-genomes (strain BCN) | ERR1308824 | ERX1609841 |

**Supplementary Table S2** – The sequences used to query for the presence of *STA1* and the 1162 bp deletion in the *STA1* promoter.

| **Name** | **Sequence** |
|---|---|
| STA1_BLAST | TTCCAACTGCACTAGTTCCTAGAGGATCCTCCTCTAGCAAC ATCACTTCCTCCGGTCCATCTTCAACTCCATTCAGCTC |
| STA1_deletion_BLAST | TGCATTTCCCAAAATTCATTCGTAGCCTTGTCAACTTAGAC TCAGTTCCACGGCGTGCAGGACGGGGTATTATGAATAAAG GATCCACGGGTAAGATTTGCTGCGCTCTCTTCTAGTTCAAG AACGGATAACTCATAGACTTACCTGTACAAGTTGTTGAAG GGTTCTCAATTGATAAAAAAGGATCTTTTGCTTCCTAA |

**Supplementary Table S3** – BLAST results for *STA1* (Genbank X02649.1). The full-length hits have been highlighted with a light grey background.

| Strain | Contig | % Identity | Alignment length | Mis-matches | Gap opens | Query start | Query end | Subject start | Subject end | e-value | bit score |
|---|---|---|---|---|---|---|---|---|---|---|---|
| **Short-read genome assemblies** | | | | | | | | | | | |
| **Strain** | **Contig** | **% Identity** | **Alignment length** | **Mis-matches** | **Gap opens** | **Query start** | **Query end** | **Subject start** | **Subject end** | **e-value** | **bit score** |
| A81062 | Scerevisiae_A62_chromosome_IX | 98.888 | 1439 | 15 | 1 | 900 | 2337 | 164632 | 166070 | 0 | 2567 |
| | Scerevisiae_A62_chromosome_IX | 97.738 | 619 | 14 | 0 | 282 | 900 | 372406 | 371788 | 0 | 1066 |
| | Scerevisiae_A62_chromosome_IX | 97.362 | 417 | 11 | 0 | 4 | 420 | 372816 | 372400 | 0 | 710 |
| WLP570 | CFF_4-19034 | 99.027 | 1439 | 13 | 1 | 900 | 2337 | 2004 | 566 | 0 | 2579 |
| | CFF_4-19035 | 98.703 | 848 | 11 | 0 | 2 | 849 | 1659 | 2506 | 0 | 1506 |
| | CFF_4-19163 | 97.9 | 619 | 13 | 0 | 282 | 900 | 20571 | 19953 | 0 | 1072 |
| OS899 | BCN_8-7617 | 99.305 | 1439 | 9 | 1 | 900 | 2337 | 5427 | 3989 | 0 | 2601 |
| | BCN_8-7599 | 100 | 686 | 0 | 0 | 179 | 864 | 1 | 686 | 0 | 1267 |
| **Long-read genome assemblies** | | | | | | | | | | | |
| **Strain** | **Contig** | **% Identity** | **Alignment length** | **Mis-matches** | **Gap opens** | **Query start** | **Query end** | **Subject start** | **Subject end** | **e-value** | **bit score** |
| A81062 | A81062_chrIX | 99.358 | 2337 | 14 | 1 | 2 | 2337 | 423717 | 421381 | 0 | 4231 |
| | A81062_chrIX | 98.888 | 1439 | 15 | 1 | 900 | 2337 | 166095 | 167533 | 0 | 2567 |
| | A81062_chrIX | 97.258 | 620 | 16 | 1 | 282 | 900 | 392537 | 391918 | 0 | 1050 |
| | A81062_chrIX | 96.172 | 209 | 8 | 0 | 212 | 420 | 392739 | 392531 | 1.92E-93 | 342 |
| WLP570 | CFF_chrX | 97.523 | 2342 | 21 | 27 | 2 | 2333 | 731041 | 728727 | 0 | 3969 |
| | CFF_chrIX | 98.75 | 1440 | 16 | 2 | 900 | 2337 | 166688 | 168127 | 0 | 2558 |
| | CFF_chrIX | 97.558 | 778 | 19 | 0 | 123 | 900 | 380269 | 379492 | 0 | 1332 |
| OS899 | BCN_chrII | 99.872 | 2338 | 2 | 1 | 1 | 2337 | 16056 | 18393 | 0 | 4300 |
| | BCN_chrIX | 98.749 | 1439 | 17 | 1 | 900 | 2337 | 166148 | 167586 | 0 | 2556 |
| | BCN_chrXIII | 99.743 | 778 | 2 | 0 | 123 | 900 | 673907 | 673130 | 0 | 1426 |
| | BCN_chrX | 99.357 | 778 | 2 | 3 | 123 | 900 | 186401 | 187175 | 0 | 1406 |
| | BCN_chrX | 74.57 | 582 | 116 | 19 | 323 | 900 | 186919 | 187472 | 1.91E-58 | 226 |
| | BCN_chrXIII | 74.394 | 578 | 116 | 18 | 323 | 896 | 673386 | 672837 | 3.19E-56 | 219 |

**Supplementary Table S4** – BLAST results for 'STA1_BLAST' (Supplementary Table S2) in the 1169 *S. cerevisiae* genome assemblies from Gallone et al. (2016) and Peter et al. (2018).

| Strain | Contig | % Identity | Alignment length | Mis-matches | Gap opens | Query start | Query end | Subject start | Subject end | e-value | bit score |
|---|---|---|---|---|---|---|---|---|---|---|---|
| AAQ | AAQ_3-7332 | 100 | 79 | 0 | 0 | 1 | 79 | 695 | 773 | 5.66E-34 | 147 |
| AAR | AAR_3-7571 | 100 | 79 | 0 | 0 | 1 | 79 | 767 | 689 | 5.66E-34 | 147 |
| AEA | AEA_8-6787 | 100 | 79 | 0 | 0 | 1 | 79 | 767 | 689 | 5.66E-34 | 147 |
| AEQ | AEQ_3-11697 | 100 | 79 | 0 | 0 | 1 | 79 | 695 | 773 | 5.66E-34 | 147 |
| AFA | AFA_4-7853 | 100 | 79 | 0 | 0 | 1 | 79 | 3661 | 3583 | 5.66E-34 | 147 |
| AFB | AFB_4-8105 | 100 | 79 | 0 | 0 | 1 | 79 | 580 | 502 | 5.66E-34 | 147 |
| AFP | AFP_1-7540 | 100 | 79 | 0 | 0 | 1 | 79 | 3659 | 3581 | 5.66E-34 | 147 |
| ALI | ALI_5-7132 | 100 | 79 | 0 | 0 | 1 | 79 | 88 | 10 | 5.66E-34 | 147 |
| AQG | AQG_4-15855 | 100 | 79 | 0 | 0 | 1 | 79 | 767 | 689 | 5.66E-34 | 147 |
| AQH | AQH_2-39734 | 100 | 79 | 0 | 0 | 1 | 79 | 958 | 1036 | 5.66E-34 | 147 |
| BCB | BCB_3-7948 | 100 | 79 | 0 | 0 | 1 | 79 | 88 | 10 | 5.66E-34 | 147 |
| BCC | BCC_3-6681 | 100 | 79 | 0 | 0 | 1 | 79 | 88 | 10 | 5.66E-34 | 147 |
| BCE | BCE_8-6067 | 100 | 79 | 0 | 0 | 1 | 79 | 32 | 110 | 5.66E-34 | 147 |
| BCF | BCF_8-6617 | 100 | 79 | 0 | 0 | 1 | 79 | 88 | 10 | 5.66E-34 | 147 |
| BCI | BCI_8-6529 | 100 | 79 | 0 | 0 | 1 | 79 | 32 | 110 | 5.66E-34 | 147 |
| BCK | BCK_8-7150 | 100 | 79 | 0 | 0 | 1 | 79 | 32 | 110 | 5.66E-34 | 147 |
| BCL | BCL_5-6462 | 100 | 79 | 0 | 0 | 1 | 79 | 32 | 110 | 5.66E-34 | 147 |
| BCN | BCN_8-4221 | 100 | 79 | 0 | 0 | 1 | 79 | 32 | 110 | 5.66E-34 | 147 |
| BCP | BCP_8-5410 | 100 | 79 | 0 | 0 | 1 | 79 | 88 | 10 | 5.66E-34 | 147 |
| BDP | BDP_5-6002 | 100 | 79 | 0 | 0 | 1 | 79 | 88 | 10 | 5.66E-34 | 147 |
| BDQ | BDQ_6-7824 | 100 | 79 | 0 | 0 | 1 | 79 | 32 | 110 | 5.66E-34 | 147 |
| BDR | BDR_6-6225 | 100 | 79 | 0 | 0 | 1 | 79 | 32 | 110 | 5.66E-34 | 147 |
| BDS | BDS_6-8316 | 100 | 79 | 0 | 0 | 1 | 79 | 41 | 119 | 5.66E-34 | 147 |
| BDT | BDT_6-11671 | 100 | 79 | 0 | 0 | 1 | 79 | 767 | 689 | 5.66E-34 | 147 |
| BEB | BEB_6_C37T3ACXX.IND41b-7767 | 100 | 79 | 0 | 0 | 1 | 79 | 88 | 10 | 5.66E-34 | 147 |

| | | | | | | | | | | | |
|---|---|---|---|---|---|---|---|---|---|---|---|
| BED | BED_6-6933 | 100 | 79 | 0 | 0 | 1 | 79 | 88 | 10 | 5.66E-34 | 147 |
| beer002 | MCAA01000038.1 | 100 | 79 | 0 | 0 | 1 | 79 | 3980 | 4058 | 5.66E-34 | 147 |
| beer004 | CM007127.1 | 100 | 79 | 0 | 0 | 1 | 79 | 398847 | 398769 | 5.66E-34 | 147 |
| beer011 | CM007015.1 | 100 | 79 | 0 | 0 | 1 | 79 | 382411 | 382333 | 5.66E-34 | 147 |
| beer013 | CM006967.1 | 100 | 79 | 0 | 0 | 1 | 79 | 396002 | 395924 | 5.66E-34 | 147 |
| beer021 | CM006855.1 | 100 | 79 | 0 | 0 | 1 | 79 | 397405 | 397327 | 5.66E-34 | 147 |
| beer032 | CM006663.1 | 100 | 79 | 0 | 0 | 1 | 79 | 395834 | 395756 | 5.66E-34 | 147 |
| beer034 | CM006647.1 | 100 | 79 | 0 | 0 | 1 | 79 | 399862 | 399784 | 5.66E-34 | 147 |
| beer039 | CM006567.1 | 100 | 79 | 0 | 0 | 1 | 79 | 392082 | 392004 | 5.66E-34 | 147 |
| beer040 | CM006551.1 | 100 | 79 | 0 | 0 | 1 | 79 | 387784 | 387706 | 5.66E-34 | 147 |
| beer059 | CM006247.1 | 100 | 79 | 0 | 0 | 1 | 79 | 186841 | 186919 | 5.66E-34 | 147 |
| beer062 | CM006199.1 | 100 | 79 | 0 | 0 | 1 | 79 | 392434 | 392356 | 5.66E-34 | 147 |
| beer080 | CM005911.1 | 100 | 79 | 0 | 0 | 1 | 79 | 408118 | 408040 | 5.66E-34 | 147 |
| beer083 | CM005863.1 | 100 | 79 | 0 | 0 | 1 | 79 | 404330 | 404252 | 5.66E-34 | 147 |
| beer084 | MBWW01000093.1 | 100 | 79 | 0 | 0 | 1 | 79 | 1143 | 1065 | 5.66E-34 | 147 |
| beer085 | CM005831.1 | 100 | 79 | 0 | 0 | 1 | 79 | 396106 | 396028 | 5.66E-34 | 147 |
| beer086 | CM005815.1 | 100 | 79 | 0 | 0 | 1 | 79 | 400774 | 400696 | 5.66E-34 | 147 |
| beer091 | CM005735.1 | 100 | 79 | 0 | 0 | 1 | 79 | 393037 | 392959 | 5.66E-34 | 147 |
| beer092 | CM005719.1 | 100 | 79 | 0 | 0 | 1 | 79 | 402761 | 402683 | 5.66E-34 | 147 |
| BMV | BMV_2-6044 | 100 | 79 | 0 | 0 | 1 | 79 | 88 | 10 | 5.66E-34 | 147 |
| BNA | BNA_4-6466 | 100 | 79 | 0 | 0 | 1 | 79 | 32 | 110 | 5.66E-34 | 147 |
| BNC | BNC_4-9323 | 100 | 79 | 0 | 0 | 1 | 79 | 88 | 10 | 5.66E-34 | 147 |
| BRM | BRM_4-7163 | 100 | 79 | 0 | 0 | 1 | 79 | 6097 | 6019 | 5.66E-34 | 147 |
| CFF | CFF_4-19035 | 100 | 79 | 0 | 0 | 1 | 79 | 1755 | 1833 | 5.66E-34 | 147 |
| CRE | CRE_2-37810 | 100 | 79 | 0 | 0 | 1 | 79 | 752 | 674 | 5.66E-34 | 147 |
| spirits007 | CM005159.1 | 100 | 79 | 0 | 0 | 1 | 79 | 392416 | 392338 | 5.66E-34 | 147 |
| spirits008 | CM005143.1 | 100 | 79 | 0 | 0 | 1 | 79 | 408577 | 408499 | 5.66E-34 | 147 |
| wine019 | CM004807.1 | 100 | 79 | 0 | 0 | 1 | 79 | 180241 | 180319 | 5.66E-34 | 147 |
| YAB | YAB-7810 | 100 | 79 | 0 | 0 | 1 | 79 | 700 | 778 | 5.66E-34 | 147 |

**Supplementary Table S5** – The assembly statistics for *S. cerevisiae* WY3711 and A81062

| | **WY3711** | **A81062** |
|---|---|---|
| Total sequence count | 17 | 18 |
| Total sequence length | 12337335 | 12525172 |
| Min sequence length | 75369 | 83792 |
| Max sequence length | 1479817 | 1507828 |
| Mean sequence length | 725726 | 695843 |
| Median sequence length | 749327 | 735604 |
| N50 | 910883 | 918893 |
| L50 | 6 | 6 |
| N90 | 448377 | 424662 |
| L90 | 13 | 13 |
| A% | 30.79 | 30.85 |
| T% | 30.73 | 30.77 |
| G% | 19.23 | 19.16 |
| C% | 19.26 | 19.22 |
| AT% | 61.52 | 61.62 |
| GC% | 38.48 | 38.38 |
| N% | 0.00 | 0.00 |

**Supplementary Table S6** – BLAST results for *MAL11*, *MAL31* and *MTT1* (GenBank LT594281) in *S. cerevisiae* WY3711. Hits from different queries with the same letter in final column (*) are the same subject sequence.

| Query | Contig | % Identity | Alignment length | Mis-matches | Gap opens | Query start | Query end | Subject start | Subject end | e-value | bit score | * |
|---|---|---|---|---|---|---|---|---|---|---|---|---|
| *MAL11* | 0 hits found | | | | | | | | | | | |
| *MAL31* | chrVII | 97.243 | 1850 | 37 | 14 | 1 | 1845 | 1094456 | 1092616 | 0 | 3121 | a |
| *MAL31* | chrII | 91.905 | 1853 | 139 | 8 | 1 | 1845 | 856240 | 854391 | 0 | 2580 | b |
| *MAL31* | chrIII | 91.707 | 1857 | 135 | 16 | 1 | 1845 | 322468 | 320619 | 0 | 2558 | c |
| *MAL31* | chrXI | 91.604 | 1858 | 137 | 16 | 1 | 1845 | 655300 | 653449 | 0 | 2549 | d |
| *MAL31* | chrII | 90.081 | 1855 | 170 | 11 | 1 | 1845 | 840081 | 838231 | 0 | 2394 | e |
| *MTT1* (LT594281) | chrII | 97.896 | 1854 | 30 | 8 | 1 | 1848 | 840081 | 838231 | 0 | 3199 | e |
| *MTT1* (LT594281) | chrIII | 95.356 | 1852 | 80 | 6 | 1 | 1848 | 322468 | 320619 | 0 | 2939 | c |
| *MTT1* (LT594281) | chrXI | 95.038 | 1854 | 84 | 8 | 1 | 1848 | 655300 | 653449 | 0 | 2907 | d |
| *MTT1* (LT594281) | chrII | 94.941 | 1858 | 76 | 15 | 1 | 1848 | 856240 | 854391 | 0 | 2894 | b |
| *MTT1* (LT594281) | chrVII | 89.32 | 1854 | 179 | 16 | 1 | 1848 | 1094456 | 1092616 | 0 | 2309 | a |
